# Profile of a flower: How rates of morphological evolution drive floral diversification in Ericales

**DOI:** 10.1101/2022.11.28.518258

**Authors:** Julian Herting, Jürg Schönenberger, Hervé Sauquet

## Abstract

**Premise of the Study:** Recent studies of floral disparity in the asterid order Ericales have shown that flowers vary strongly among families and that disparity is unequally distributed between the three flower modules (perianth, androecium, gynoecium). However, it remains unknown whether these patterns are driven by heterogeneous rates of morphological evolution or other factors.

**Methods:** Here, we compiled a dataset of 33 floral characters scored for 414 extant ericalean species sampled from 346 genera and all 22 families. We conducted ancestral state reconstructions using an equal rates Markov models for each trait. We used the rates estimated during the ancestral state reconstruction for comparing evolutionary rates between flower modules, creating a “rate profile” of ericalean flowers.

**Key Results:** The androecium exhibits the highest evolutionary rates across most characters, whereas most perianth and gynoecium characters evolve slower. High and low rates of morphological evolution can result in high floral disparity in Ericales. Analyses of an angiosperm-wide floral dataset reveal that this pattern appears to be conserved across most major angiosperm clades.

**Conclusions:** Elevated rates of morphological evolution in the androecium of Ericales may explain the higher disparity reported for this floral module. We discuss the implications of heterogenous morphological rates of evolution among floral modules from a functional perspective. Comparing rates of morphological evolution through rate profiles proves to be a powerful tool in understanding floral evolution.

## INTRODUCTION

The asterid order Ericales comprises 22 families recognised by APG IV (Angiosperm Phylogeny Group, 2016) with 346 genera and about 12,000 species (Stevens, 2017). In earlier, pre-molecular classification systems, the families now recognised as part of Ericales were scattered in many different orders (e.g., Cronquist 1981). This was largely due to high levels of floral diversity and disparity exhibited by Ericales (Figure 1). Flowers in Ericales may be zygomorphic (e.g., Balsaminaceae; Figure 1I) or actinomorphic (e.g., Actinidiaceae; Figure 1F), show various forms of polystemony (e.g., Lecythidaceae or Actinidiaceae; Figure 1D, F), or varying degrees of fusion between perianth parts (Schönenberger and Grenhagen, 2005; Schönenberger et al., 2010; von Balthazar and Schönenberger, 2013; Zhang and Schönenberger, 2014; Löfstrand and Schönenberger, 2015b; a; Löfstrand et al., 2016). Although the monophyly of Ericales is well-supported by numerous studies (Bremer et al., 2002; Soltis et al., 2011; One Thousand Plant Transcriptomes Initiative, 2019), the interfamilial relationships within the order have remained difficult to resolve(Anderberg et al., 2002; Schönenberger et al., 2005). Recent phylogenetic studies based on Sanger sequencing (Rose et al., 2018) and nuclear genomic data (Larson et al., 2020) studies have confirmed supra-familial clades initially proposed by Schönenberger et al. (2005). Both studies agree that backbone relationships between these clades and the remaining families have low support and short branch lengths, hinting at a rapid diversification of Ericales in the Late Cretaceous (Schönenberger et al., 2005; Rose et al., 2018; Larson et al., 2020).

**Figure 1:**
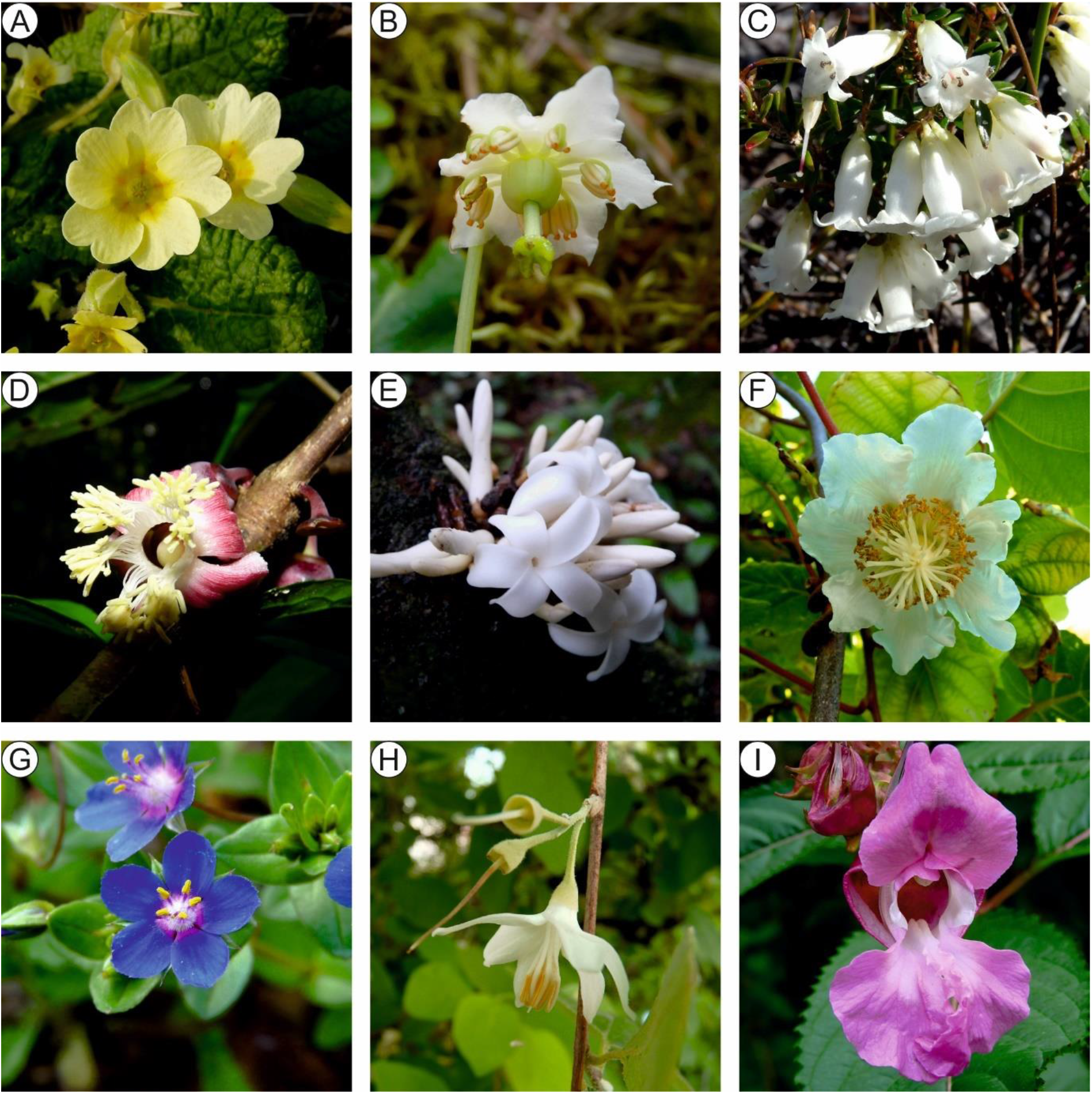
Floral diversity in Ericales. (A) *Primula elatior* (Primulaceae); (B) *Moneses uniflora* (Ericaceae); (C) *Epacris obtusifolia* (Ericaceae); (D) *Brazzeia* sp. (Lecythidaceae); (E) *Diospyros fragrans* (Ebenaceae); (F) *Actinidia chinensis* (Actinidiaceae); (G) *Lysimachia foemina* (Primulaceae); (H) *Styrax officinalis* (Styracaceae); (I) *Impatiens glandulifera* (Balsaminaceae). Photos by ©Hervé Sauquet.

Flowers are generally divided into three functional modules: the perianth, the androecium, and the gynoecium. The perianth usually serves as protection for the flower (usually the calyx) and as attraction for animal pollinators (usually the corolla), the androecium produces and presents pollen and may also be involved in pollinator attraction, and lastly the gynoecium produces ovules, seeds after fertilisation, as well as the fruit, and disseminates the seeds (Endress, 1996, 2011). Comparing pairwise dissimilarity and the floral morphospace of 30 floral traits, Chartier et al. (2017) showed that floral disparity among Ericales varies amongfamilies and flower modules. Disparity was found to be highest in the androecium and lowest in the perianth. This was attributed to different selective regimes, varying degrees of module complexity, and developmental constraints influencing the flower modules in different ways. For example, multiple origins of polystemony and the apparent lability of androecium traits have been proposed as the main drivers of the high disparity found in the androecium (Schönenberger et al., 2005; Endress, 2011; Löfstrand et al., 2016; Chartier et al., 2017). Four families (Ericaceae, Lecythidaceae, Primulaceae, and Sapotaceae) were found to account for more than half of the disparity among Ericalean flowers (Chartier et al., 2017). Disparity measured as pairwise dissimilarity does not take phylogeny into account making the results highly dependent on the taxonomic scale (Foote, 1999). Thus, using evolutionary rates, which consider phylogeny as a measurement of variation might lead to more insights. However, it has not yet been studied how disparity and evolutionary rates interact, the obvious assumptions being that the faster a trait evolved, the higher its disparity, or vice versa.

Rates of morphological evolution (henceforth morphological rates), typically measured as the number of transitions of a trait in a million years, have been a central parameter of important macroevolutionary models for several decades (e.g., Simpson 1944; Pagel 1994; Lee et al. 2013). Morphological rates are mainly used, and are an important parameter, in the reconstruction of phylogenies using morphological or total evidence datasets (e.g. Lewis, 2001; Ronquist et al., 2012; Herrera et al., 2020), reconstructing ancestral states (e.g., Landis et al., 2018; Carrive et al., 2020), and to estimate divergence times in a total-evidence dating framework (e.g., Ronquist et al. 2012; Lee et al. 2014; Gavryushkina et al. 2017). A smaller number of studies have focused on morphological rates themselves as the main subject. For instance, recent work has focused on the adequacy of morphological rate models in portraying morphological evolution, and on the mismatch between models of morphological and molecular evolution, often in the context of phylogenetic reconstructions and total-evidence dating (e.g., Thomas et al. 2010; Goloboff et al. 2019; Asar et al. preprint 2022; Klopfstein et al. preprint 2020). In the same context, morphological rate heterogeneity has begun to be quantified and incorporated into models to address among-lineage heterogeneity in phylogenetic models and ancestral state reconstructions (O’Meara et al., 2006; Beaulieu et al., 2012), most often for continuous traits, but also for discrete characters (Beaulieu et al., 2013; Boyko and Beaulieu, 2021). Reyes et al. (2018) showed that morphological rates and among-lineage heterogeneity differ among floral traits, but studies of among-trait heterogeneity of morphological rates remain scarce. Heterogenous rates among floral traits immediately lead to questions about the influence of functional morphology on morphological rates. Hypothetically, traits that are associated with pollinator attraction (e.g., display size) are expected to evolve faster than traits associated with floral organization (e.g., phyllotaxis).

Here, we address these questions by quantifying rates of floral evolution in Ericales. As outlined above, the order is a well-suited clade to study the influence of among-character rate heterogeneity on flower disparity and floral evolution thanks to its remarkably diverse flowers (Figure 1) and well-studied floral morphology. We estimate rates of morphological evolution for 32 floral traits of Ericales, compare rates among them and among floral modules, and reconstruct the ancestral state for each trait. Thus, we create a “rate profile” of the ericalean flower and discuss whether the rate of morphological evolution matches the observed floral disparity described for Ericales. Additionally, we discuss the reconstructed ancestral states for some of the traits in our dataset considering the interaction between morphological rates, floral disparity, and functional significance. Lastly, we discuss possible implications of using this approach on datasets spanning greater phylogenetic diversity by applying the same methodology to an angiosperm-wide dataset.

## METHODS

All analyses were conducted in R (R Core Team 2022) with the *tidyverse* package (Wickham et al., 2019). Further, we used the packages *ape* (Paradis and Schliep, 2019), *corHMM* (Beaulieu et al., 2013), and *phytools* (Revell, 2012) for functions specific to comparative phylogenetics.

### Morphological dataset and taxon sampling

We based taxon sampling for morphological data on the taxa present in the phylogenetic analyses by Rose et al. (2018) and taxa used in the analyses of floral disparity by Chartier et al. (2017), to maximise overlap between the two datasets. To fully represent phylogenetic diversity at the generic level, we included at least one species per genus. Our dataset comprises morphological data for 414 species sampled from all 346 Ericalean genera and all 22 families (Appendix S1; see Supplemental Data with this article). Morphological data for all species used in the present analyses were recorded in the PROTEUS database (Sauquet, 2019). We chose to analyse 32 floral characters (four general, eleven perianth, ten androecium, and seven gynoecium traits) (see Appendix S1). These characters were chosen from the 37 characters used to analyse floral disparity by Chartier et al. (2017). Data for 286 species had already been recorded in PROTEUS by Chartier et al. (2017). The remaining 128 species were recorded for this study. All taxa were scored from published literature. The complete list of 9964 data records (each linked to an explicit source) and 431 references used is provided as supplementary material Appendix S1.

### Morphological rate estimation

We used the ancestral state reconstruction function corHMM implemented in *corHMM* to estimate morphological rates (transition rates). Ancestral state reconstructions and therefore morphological rate estimation were inferred on the dated phylogeny reconstructed by Rose et al. (2018). We pruned the dated phylogeny to the 414 taxa represented in the morphological dataset. We ran the analyses for each character independently and forcing equal transition rates (“ER” option in *corHMM*). We elected to only use the equal rates model for several reasons. In all rates different (ARD) models more parameters need to be estimated, introducing more uncertainty. ARD models face several challenges which make them ill-fitting for the analyses we conduct here; they are prone to artefacts, such as overestimation of rates if origins of a state are located on a short terminal branch (Goldberg and Igić, 2008; Sauquet et al., 2017; Reyes et al., 2018). Additionally, using the ER model enhances comparability between rates among traits. We then extracted the morphological rates from the resulting transition matrices of the equal rates analyses. We plotted the resulting likelihoods of the reconstruction on a randomly drawn stochastic character map simulated using *corHMM* (Beaulieu et al., 2013). The extracted rates were then used to compare the tempo of evolution between characters and floral modules. We performed an ANOVA to test whether the mean morphological rates differed significantly among flower modules.

To estimate the absolute number of back-and-forth transitions between states we also reconstructed 1000 stochastic maps (makeSimmap function in *corHMM*) for each character. We then extracted the number of transitions from each and each estimated stochastic map individually trait using the density.multisimmap function of *phytools*, resulting in two transition numbers for binary traits and six for traits with three states for each map. We proceeded to calculate the mean and 95% credibility intervals for each trait (see Appendix S2).

### Angiosperm-wide analysis

For our angiosperm-wide analyses, we used the morphological dataset of Schönenberger et al. (2020) comprising 792 species sampled from 63 orders and 372 families of angiosperms, which represented an updated and more complete version of the eFLOWER dataset originally presented by Sauquet et al. (2017). We used the time-calibrated tree of Magallón et al. (2015) as the base of inference. We applied the same methodology of estimating morphological rates for the angiosperm-wide dataset as was used for the Ericales dataset. If a trait was uniform in the analysed clade (i.e., all tips have the same state), the trait was excluded from the analyses for this clade because it is not possible to estimate morphological rates for uniform traits. Similarly, we excluded a trait from plotting in the bar plots if its morphological rate was so high that is could be considered an outlier due to artifacts linked with rare autapomorphic states at this scale (e.g., connective extension in Malvidae with a rate > 1 t/ma).

## RESULTS

### Evolutionary rates of Ericales flower modules

Rates of morphological evolution vary dramatically among flower modules and among traits of the same module. These differences in morphological rates become apparent in the rate profile of ericalean floral evolution, which shows that most androecium traits evolve faster than perianth or gynoecium traits (Figure 2).

**Figure 2:**
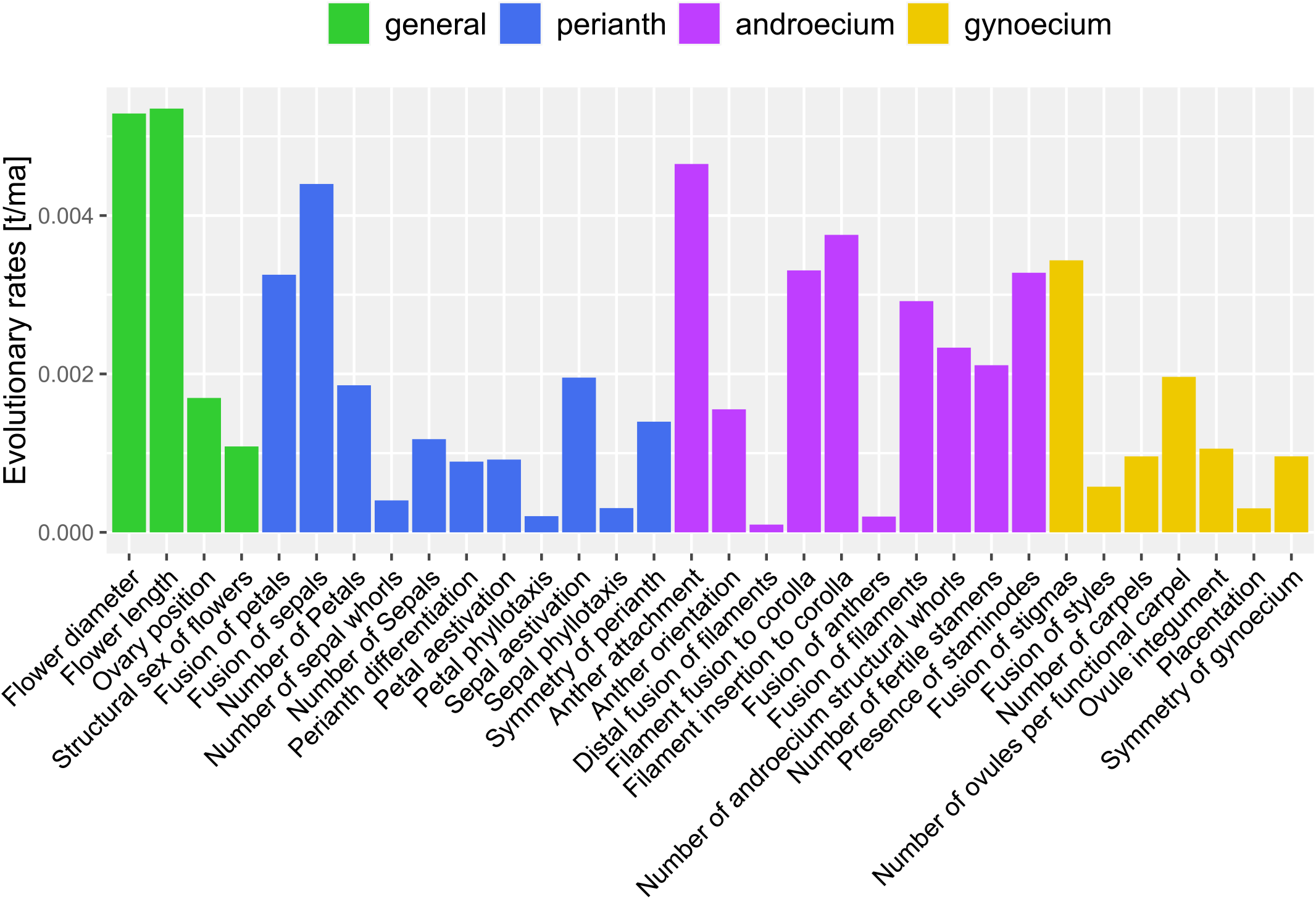
Rate profile of Ericales estimated using an equal rates Markov model. Each trait was analysed independently. Traits are shown here grouped by the flower module it belongs to. Most androecium traits evolve at rates higher than 0.002 t/ma, whereas most traits of the other modules evolve at a slower rate.

The fastest evolving traits are the two traits depicting flower size (Flower length and Flower diameter) with morphological rates above 0.005 transitions per million years (henceforth t/ma). Of the three flower modules the androecium has the fastest evolving traits compared to the other flower modules (Figure 2, 3). Androecium traits have a median morphological rate of 0.0026 t/ma (Figure 3), with only two of ten trait showing rates below 0.001 t/ma (Figure 2). The maximum rate for that module is 0.004 t/ma estimated for the anther attachment character, while the slowest evolving androecium character is distal fusion of filaments at a rate of 0.000097 t/ma (Figure 2A). Perianth traits evolve slower than androecium traits, but faster than gynoecium traits on average having a median morphological rate of 0.0011 t/ma (Figure 3). The fusion of sepals, i.e. whether sepals are laterally fused into a tube, evolves the fastest among perianth traits (0.0043 t/ma), whereas petal phyllotaxis evolves the slowest (0.0002 t/ma) (Figure 2A). The median rate of morphological evolution in the gynoecium is below 0.001 at 0.00095 t/ma and thus the lowest of the three modules (Figure 3). Among the gynoecium traits, placentation evolves the slowest (0.0003 t/ma) and the fusion of stigmas evolves the fastest (0.0034 t/ma) (Figure 2). However, despite this apparent pattern of morphological rates differing among flower modules, we infer no significant difference between the morphological rates (p-value ANOVA = 0.17).

**Figure 3:**
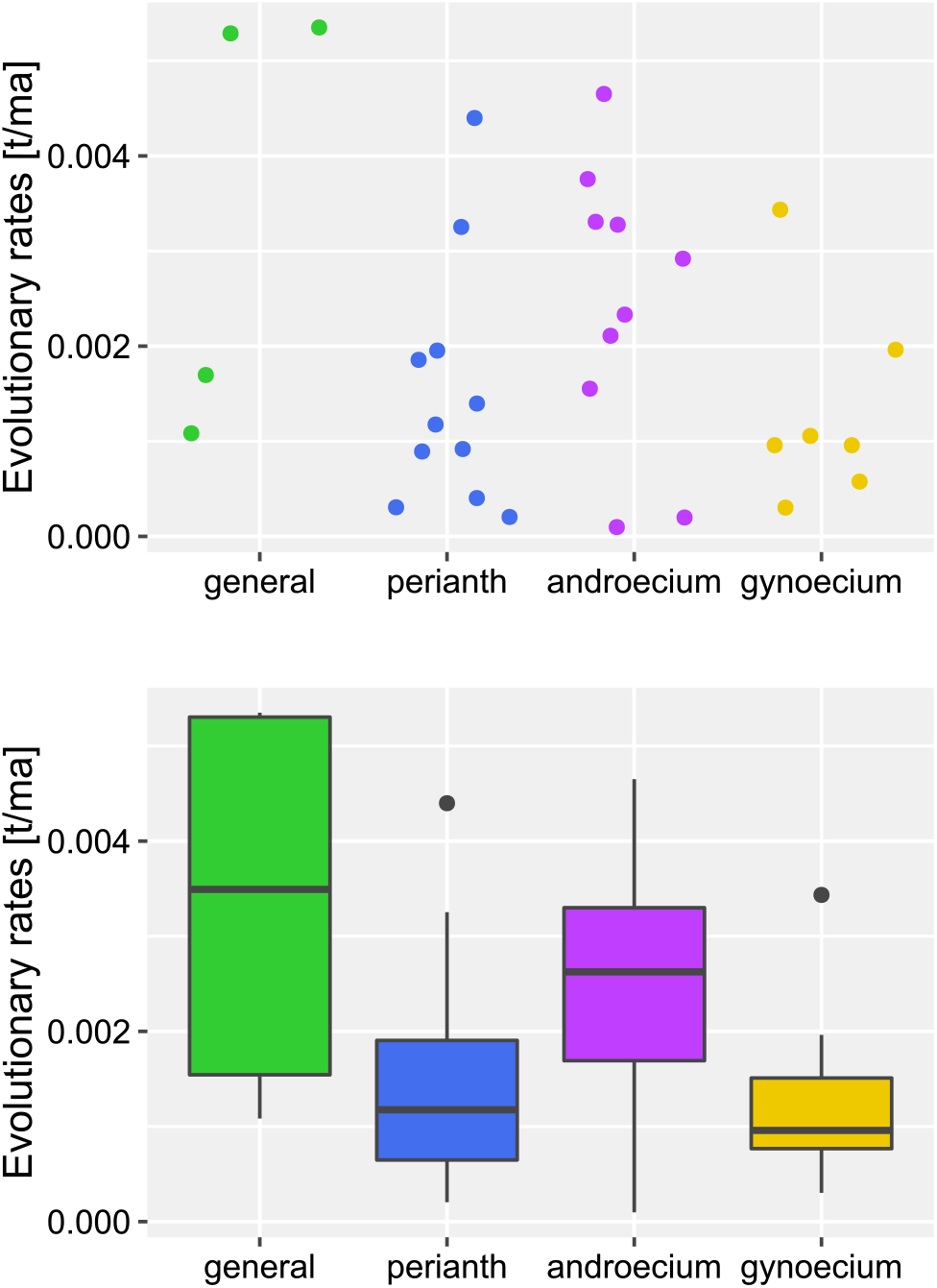
Scatter- and boxplot of morphological rates summarised for the three flower modules and general traits. The median morphological rate of the androecium is markedly higher than for the other two modules.

### Angiosperm-wide morphological rate patterns

Morphological rates inferred on the angiosperm-wide dataset, for 15 major clades, show a similar pattern or rate profile as the focussed analyses of the Ericales dataset (Figure 3,4). The rate profiles of all major clades show that androecium traits have the highest morphological rates, and that the androecium consistently includes most traits with high rates (Figure 4, for larger version with trait names see Appendix S3). Magnoliidae are the only clear exception to this pattern. In their rate profile, morphological rates of gynoecium traits are elevated compared to other major clades. Additionally, gynoecium traits are the fastest evolving traits of Magnoliidae, making them an exception compared to the other major clades (Figure 4). In absolute values, rates for this dataset seldom exceed 0.01 t/ma.

**Figure 4:**
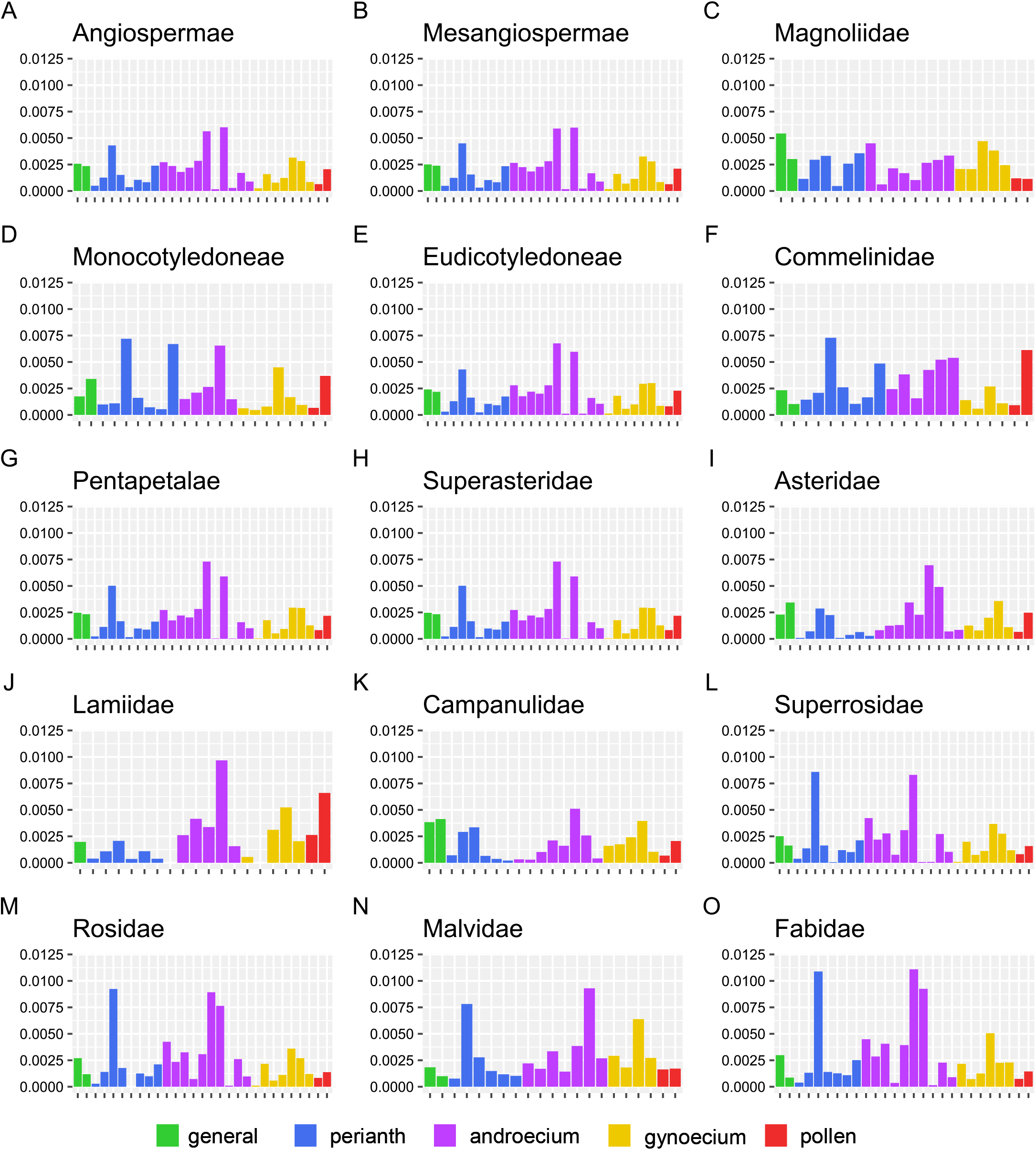
Rate profiles of major angiosperm clades. In all major angiosperm clades, except Magnoliidae, the androecium exhibits the most traits with high morphological rates. (A) all angiosperms; (B) Mesangiospermae; (C) Magnoliidae; (D) Monocotyledoneae; (E) Eudicotyledoneae; (F) Commelinidae; (G) Pentapetalae; (H) Superasteridae; (I) Asteridae; (J) Lamiidae; (K) Campanulidae; (L) Superrosidae; (M) Rosidae; (N) Malvidae; (O) Fabidae. Green: general traits; blue: perianth; purple: androecium; yellow: gynoecium; red: pollen.

## DISCUSSION

### Morphological rates and floral disparity

Our analyses show that morphological rates differ among flower modules, with the androecium traits evolving faster than traits of the other two modules (Figure 2,3). However, fast and slow evolving traits can be found in all three modules (Figure 2). Comparatively faster morphological rates for the androecium match the pattern of floral disparity found in Ericales (Chartier et al. 2017). Floral disparity, measured as pairwise dissimilarity, is higher in the androecium compared to other floral modules (Chartier et al. 2017). However, comparison between our analyses and the analyses of Chartier et al. (2017) reveal that, although the overall patterns of evolutionary rate and floral disparity match among modules, this is not always the case at the level of individual traits. High levels of disparity may result from both fast and slow morphological rates, with the rate affecting the distribution of the different states on the phylogeny.

For example, anther orientation and integument number are highly disparate among Ericales (Chartier et al. 2017 Figure S2), but according to our analyses they have low to medium morphological rates compared with other traits (Figure 2). On the other hand, there are traits that exhibit both high disparity in Ericales and a high rate of morphological evolution, for example fusion of the filament to the corolla (Figure 2) (Chartier et al. 2017 Figure S2). In the former case of high disparity paired with low morphological rates, stochastic mapping shows 6-11 (95% HPD) transitions from the ancestral bitegmic ovule to the unitegmic ovule and 1-8 (95% HPD) transitions back from unitegmic to bitegmic (Appendix S2, S4). However, there is a marked difference on where in the phylogeny these transitions occur, transitions to unitegmy tend to occur at the base of family clades (e.g., Polemoniaceae) or supra familial clades (e.g., clade comprising styracoids, sarracenioids and ericoids). Conversely, transitions back to bitegmy tend to occur near the tips (e.g., *Styrax*) (Appendix S2, S4). These early transitions to unitegmy result in more than half of the ericalean species analysed here being unitegmic and high disparity that is largely distributed between few clades that are homogenous for either state. Thus, few transitions of a trait early during the divergence of Ericales, with few or no transitions back (i.e. low morphological rates) result in high disparity that stems from differences among larger clades or lineages.

In contrast, high disparity that is co-occurring with high morphological rates stems from more frequent transitions between trait states within families or transitions back to the ancestral state. For example, the ancestral state reconstruction for the trait “filament fusion to corolla” shows that fusion between the filament and the petals evolved independently in at least seven larger ericalean clades and several individual species (19-30 transitions to fused, Appendix S2; Figure 5), with several species transitioning back to unfused filaments and petals (10-21 transitions back to unfused, Appendix S2; Figure 5). A similar pattern can be observed in the reconstruction of the anther attachment (Appendix S2, S4). Thus, higher rates of morphological evolution shift the disparity from differences among families and supra familial clades to mostly genera and species within a family being disparate. By virtue of its methodology, disparity measured as mean pairwise dissimilarity is agnostic to the phylogeny of the compared taxa (Foote, 1999). Therefore, we advocate that in future studies of disparity, morphological rates and ancestral states should be studied together because the latter give vital information on the distribution of disparity through time that can be further enhanced by the incorporation of fossil taxa.

**Figure 5:**
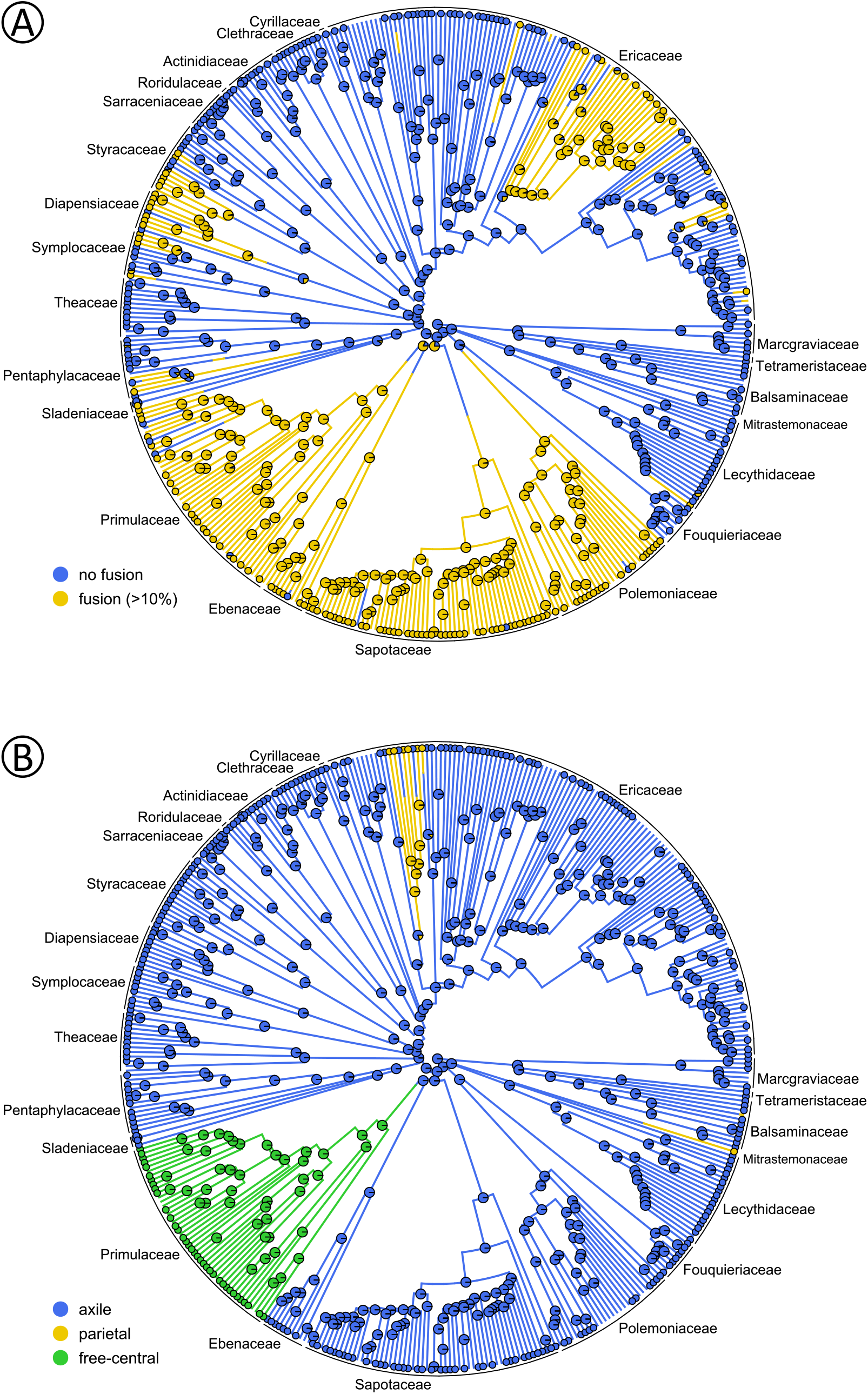
Ancestral state reconstructions using an equal rates model for: (A) the filament fusion to the corolla, blue: no fusion, yellow: more than 10% fusion; (B) the placentation, blue: axile, yellow: parietal, green: free-central.

### Rate heterogeneity among floral traits

In Ericales, rates of morphological evolution vary within two orders of magnitude among different traits. It would be expected that traits which are synapomorphic for a subclade of the clade under investigation have a comparatively low morphological rate because the new state evolved only once with no reversals (in the extreme case). An example from our dataset illustrating this, is the distal fusion of filaments, which is synapomorphic for Balsaminaceae (Schönenberger et al., 2010; von Balthazar and Schönenberger, 2013) and has the lowest morphological rate of all traits (0.00009 t/ma, Appendix S2, S4). Yet, because the distal fusion of filaments is synapomorphic for Balsaminaceae, the transition to distally fused filaments sits on a comparatively deep node of the phylogeny.

Although Ericales are known for comprising several polystemonous lineages (e.g., Lecythidaceae, Theaceae, and Actinidiaceae), the evolutionary rates of the traits facilitating polystemony (number of fertile stamens and number of androecium structural whorls) are in the medium range compared to other androecium traits, but high compared with traits of other modules (Figure 2). Polystemony in Ericales can occur in different ways, e.g., by possessing more than two whorls of stamens [e.g., in Lecythidaceae (Endress, 1996)], by producing organ pairs in part of the stamen positions [e.g., in Fouquieriaceae (Schönenberger and Grenhagen 2005)], or by expressing multiple stamens on a ring primordium [e.g., in Actinidiaceae (Löfstrand and Schönenberger, 2015a) and Pentaphylacaceae (Zhang and Schönenberger, 2014)]. Furthermore, Löfstrand and Schönenberger (2015b) conducted ancestral state reconstruction of the number of androecium whorls and conclude that both gains and losses of a second, antepetalous whorl occurred repeatedly in Ericales. Here we recover a similar pattern the of frequent transitions between one and two (and rarely three) androecium whorls (Appendix S2, S4), thus plasticity increases the morphological rate.

The two fastest evolving traits in Ericales both relate to floral size (flower length and flower diameter) with flowers of all three size categories appearing in the same clades with no clear phylogenetic pattern and no unequivocally reconstructed ancestral state (Appendix S4). For example, all three size categories are present in Sapotaceae, Ericaceae, and Lecythidaceae for both flower length and flower diameter. Traits like floral size or fusion of organs are considered traits of floral mode [sensu Endress, (1996)] which are more directly involved in the interaction between flower and pollinator and are therefore expected to be highly variable, even among species of the same genus. Differing pollination syndromes are linked to different length ratios among floral organs, for example wind-pollinated flowers have short petals in relation to the style and an extruding style (e.g. *Erica scoparia*), whereas flowers pollinated by long-proboscis flies tend to have long fused petals with a narrow aperture that restricts access, and the style only slightly extrudes from the flower tube (e.g., *Erica ventricosa*) (Reich et al., 2020). This illustrates that closely related species of *Erica* have quickly evolved different pollination syndromes by modifying floral mode traits, but the general bauplan stays the same, for example all studied flowers are tetramerous and whorled (Reich et al., 2020).

Phyllotaxis of petals and sepals are among the slowest evolving traits in Ericales (Figure 2), with only a few taxa in Theaceae having spirally arranged petals and sepals (Appendix S4). Phyllotaxis is a fundamental trait of floral organization and shifts in phyllotaxis would heavily impact other floral traits (Endress, 2006, 2011). Thus, for phyllotaxis, a low morphological rate is expected in a derived group of angiosperms such as Ericales because of the high level of synorganization among floral organs as expressed, for example, in sympetally, the fusion between filaments and corolla, and zygomorphy. Transitions away from a whorled phyllotaxis would disrupt this synorganization and may lead to decreased fitness. Therefore, high morphological rates of these traits and thereby also their high disparity can be the result of species adapting to their respective ecological niche, whereas low morphological rates may be indicative of the trait being fundamentally important for the function of the flower.

### Angiosperm-wide rate profile

The inferred rate profiles of major angiosperm clades show that the trend uncovered for Ericales (i.e., highest morphological rates in the androecium) seems to be a general pattern applicable to most major angiosperm clades. This is especially interesting because we used a different set of traits (taken from Schönenberger et al., 2020) in these analyses. Some traits are unique to each set, while 13 were used in both analyses (e.g., structural sex of the flower).

Surprisingly, the only clade in which the androecium does not evolve faster than the other floral modules is the Magnoliidae. Here the fastest evolving traits are gynoecium traits and the structural sex of the flower (Figure 4, Appendix S4). One reason for the comparatively high morphological rates of the magnoliid gynoecium may be that Magnoliidae are either multi- or unicarpellate with carpel numbers ranging from one (e.g., Laurales) to many (e.g., Magnoliaceae) (Endress, 1990, 2014), whereas the other mesangiosperm lineages in this dataset usually have two to five carpels. The comparatively low rates of morphological evolution across all flower modules in Magnoliidae are in stark contrast to them being perceived as unusually variable and evolutionarily old (Massoni et al., 2015), which further supports that high disparity may arise from slow morphological rates, as shown above in Ericales.

However, the results of this analysis must be cautiously interpreted. Although the overall sample size of 792 angiosperm taxa is among the highest for an angiosperm-wide morphological dataset so far, sample size is still limited for some major clades (e.g., 42 species of Magnoliidae). Nonetheless, we think the analysis of the angiosperm-wide dataset highlights the differences and similarities between the major clades and might serve as a starting point for future research.

### CONCLUSIONS

Rates of morphological evolution in Ericales differ among floral traits and flower modules. A similar pattern of heterogeneity among morphological rates is observed across angiosperms and Ericales, suggesting common trends of floral morphological evolution which seems to act on similar traits in different lineages. Traits that are directly associated with pollinator interaction (floral mode) tend to evolve faster, while traits of floral organisation tend to evolve slower. Both slow and fast evolving traits may be highly disparate, but the slower the morphological rate, the earlier in the evolutionary history transitions occur, resulting in large clades tending to be homogenous for either state of a trait. As shown here, discussing floral disparity in the light of morphological rates allows for a more detailed interpretation of the origin of floral diversity. Thus, we argue that rates of morphological evolution are a powerful comparative tool in studying the evolution of flowers, not only when used to compare lineages, as mostly done, but also when comparing traits and groups of traits in a rate profile as done here.

## ACKNOWLEDGEMENTS

We thank Marion Chartier, Florian Jabbour, Stefan Löfstrand, and Maria von Balthazar for contributing morphological data to the original version of Ericales dataset, which was expanded for this study. JH thanks Ruby Stephens, Yasmin Asar and other members of the Sauquet lab for insightful discussions and comments. JH was funded by the German research foundation (DFG, grant number: 491286143).

## AUTHOR CONTRIBUTIONS

J.H., J.S. and H.S. designed the study. J.H. scored new floral data for Ericales. J.H. and H.S. coordinated and curated the dataset. J.H. conducted all analyses and wrote the first draft of the paper, with subsequent contributions from all authors.

## DATA AVAILABILITY

All data records from PROTEUS used in this paper are provided as Appendix S4.

## SUPPORTING INFORMATION

Additional supporting information may be found online in the Supporting Information section at the end of the article.

***Appendix S1*** Appendix S1 contains the full list of referenced data records scored in PROTEUS.

***Appendix S2*** Appendix S2 contains supplementary results, a table listing transitions rates and number of transitions for all traits.

***Appendix S3*** Appendix S3 contains supplementary figures comprising larger versions of the flower profiles of major angiosperm clades and ancestral state reconstruction for all 32 traits analysed here.

***Appendix S4*** Appendix S4 contains supplementary figures of ancestral state reconstruction for all 32 traits analysed here.

