## Supplementary figures and images for "Profile of a flower: How rates of morphological evolution drive floral diversification in Ericales"

### Appendix S3

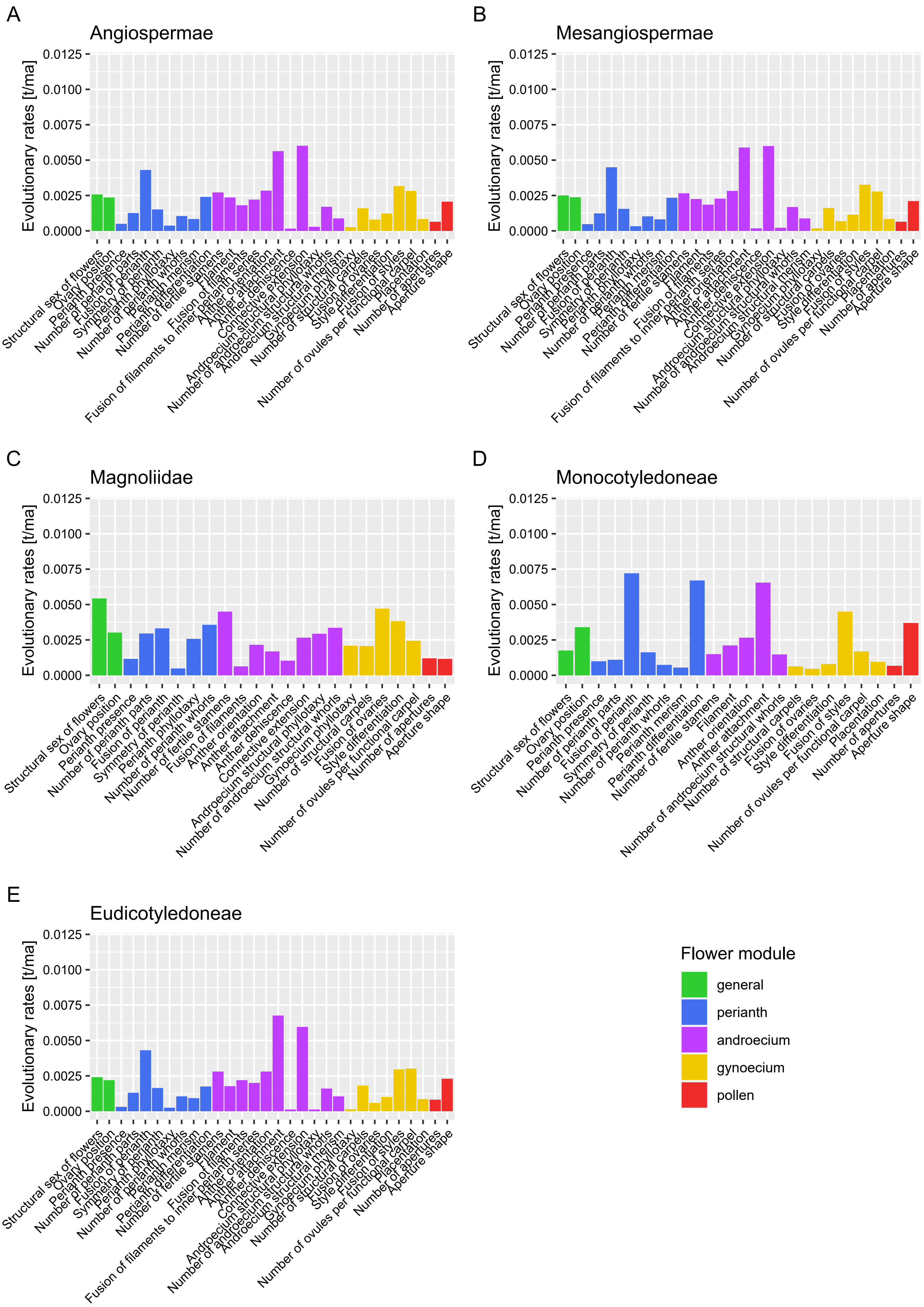

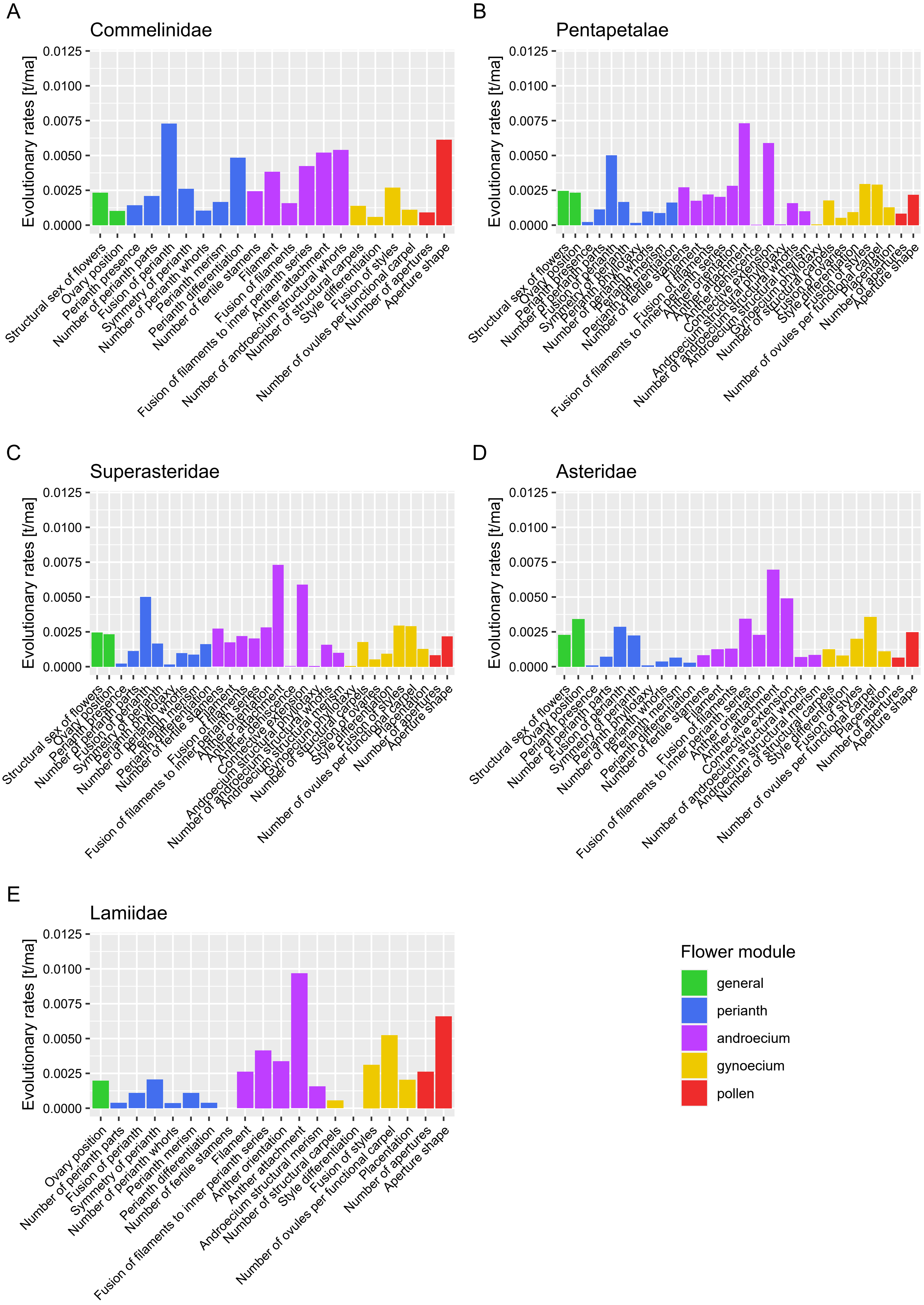

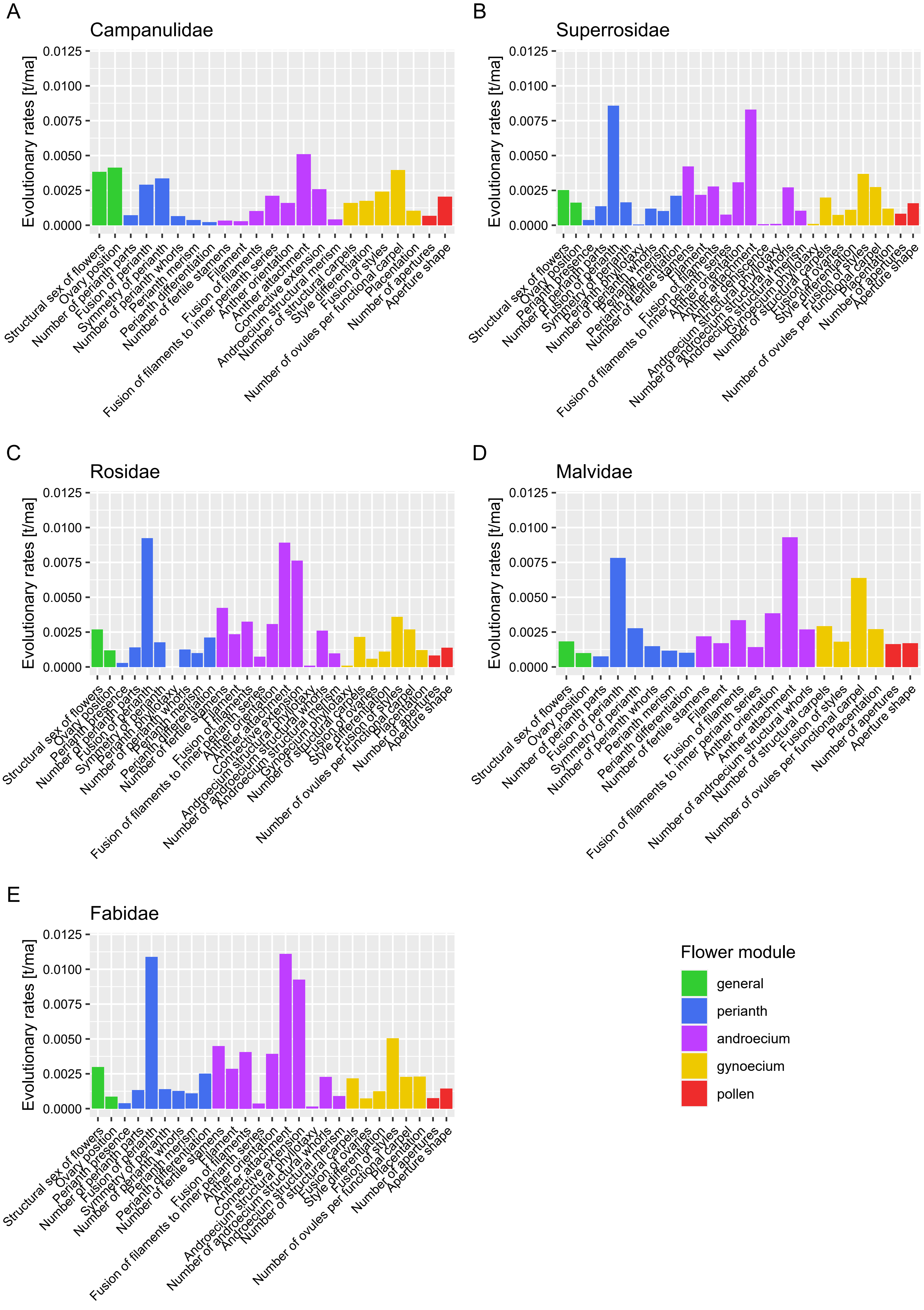
