## Appendix S4 for "Profile of a flower: How rates of morphological evolution drive floral diversification in Ericales"

### Fusion of styles

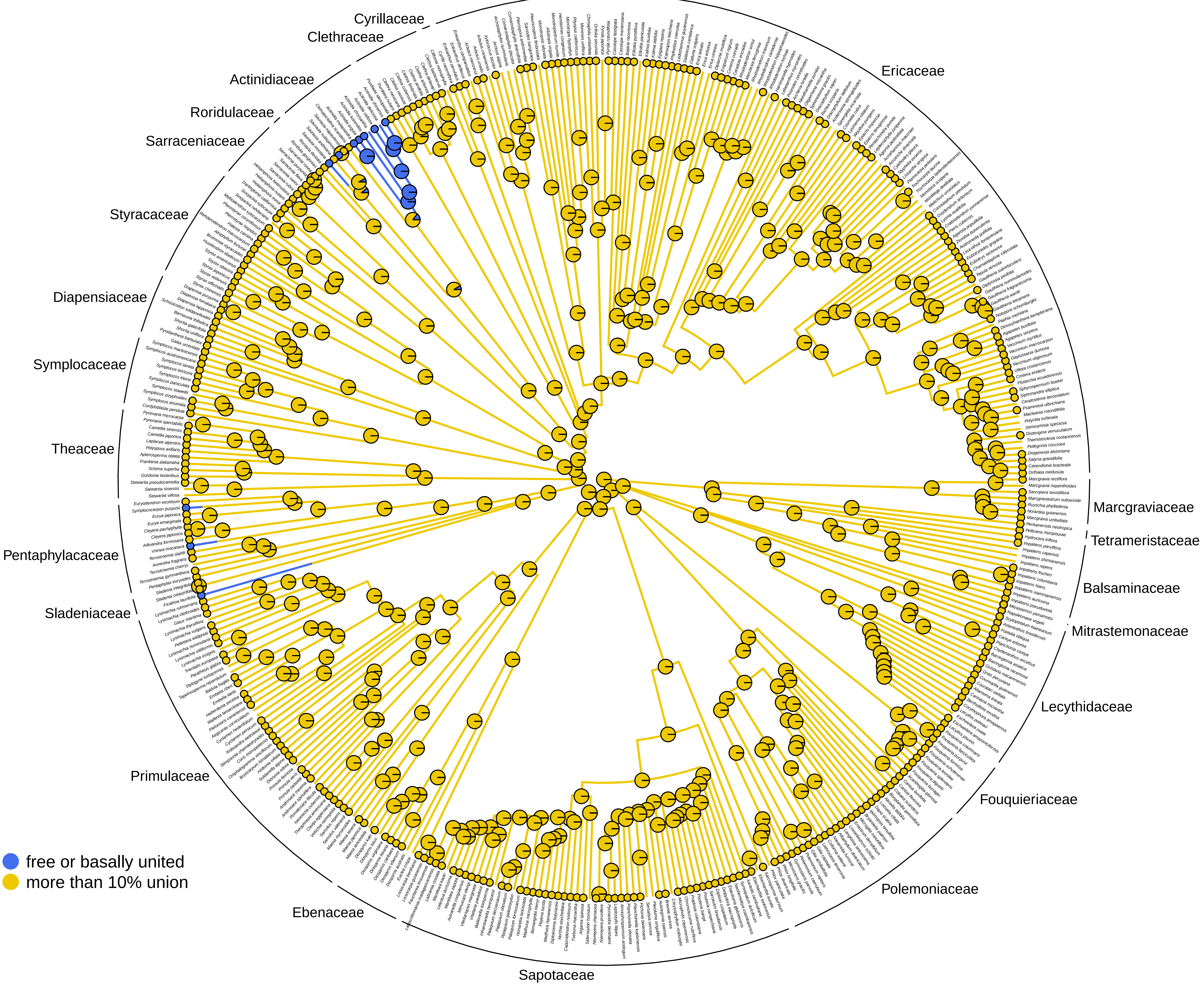

### Number of androecium structural whorls

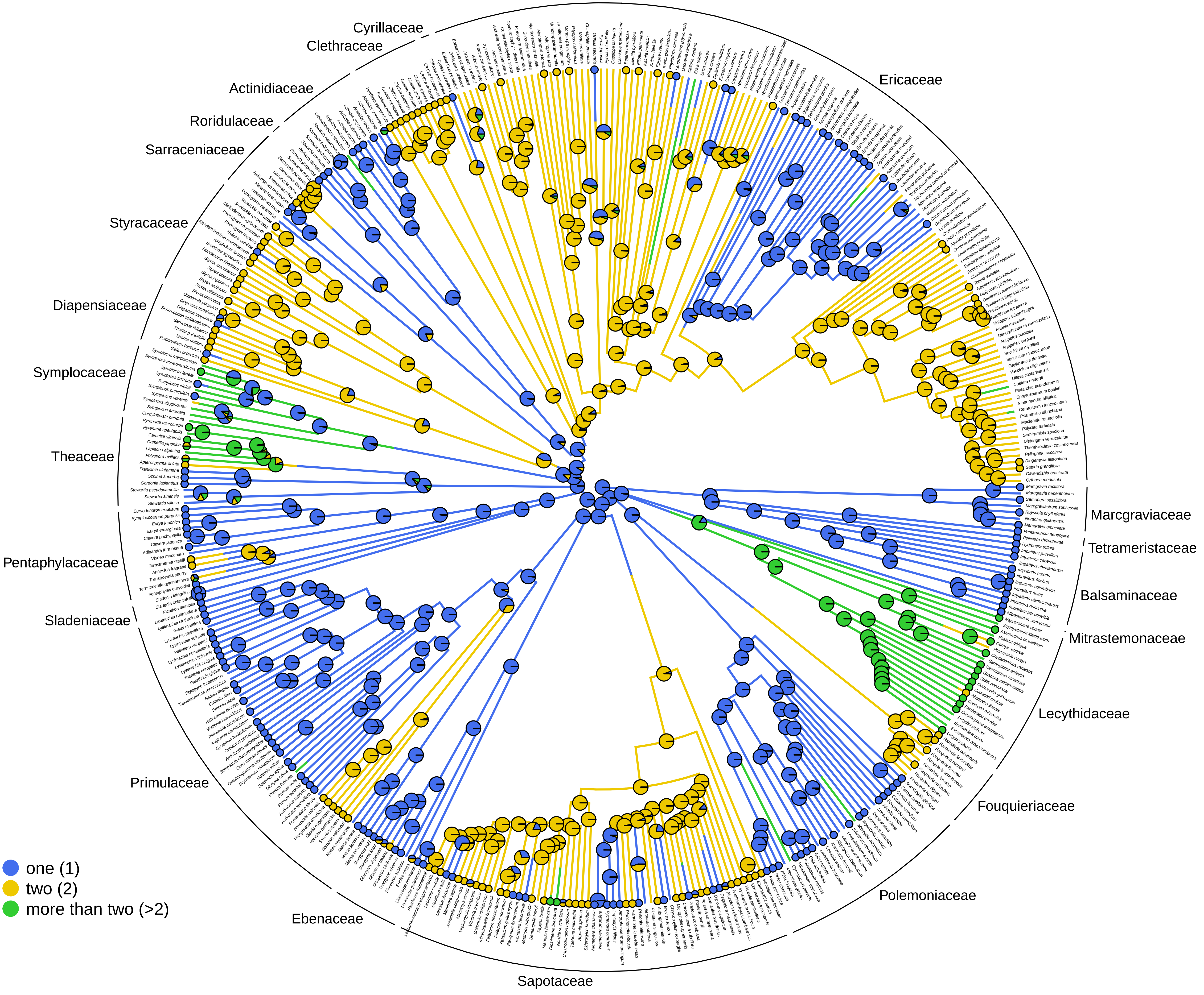

#### Number of carpels

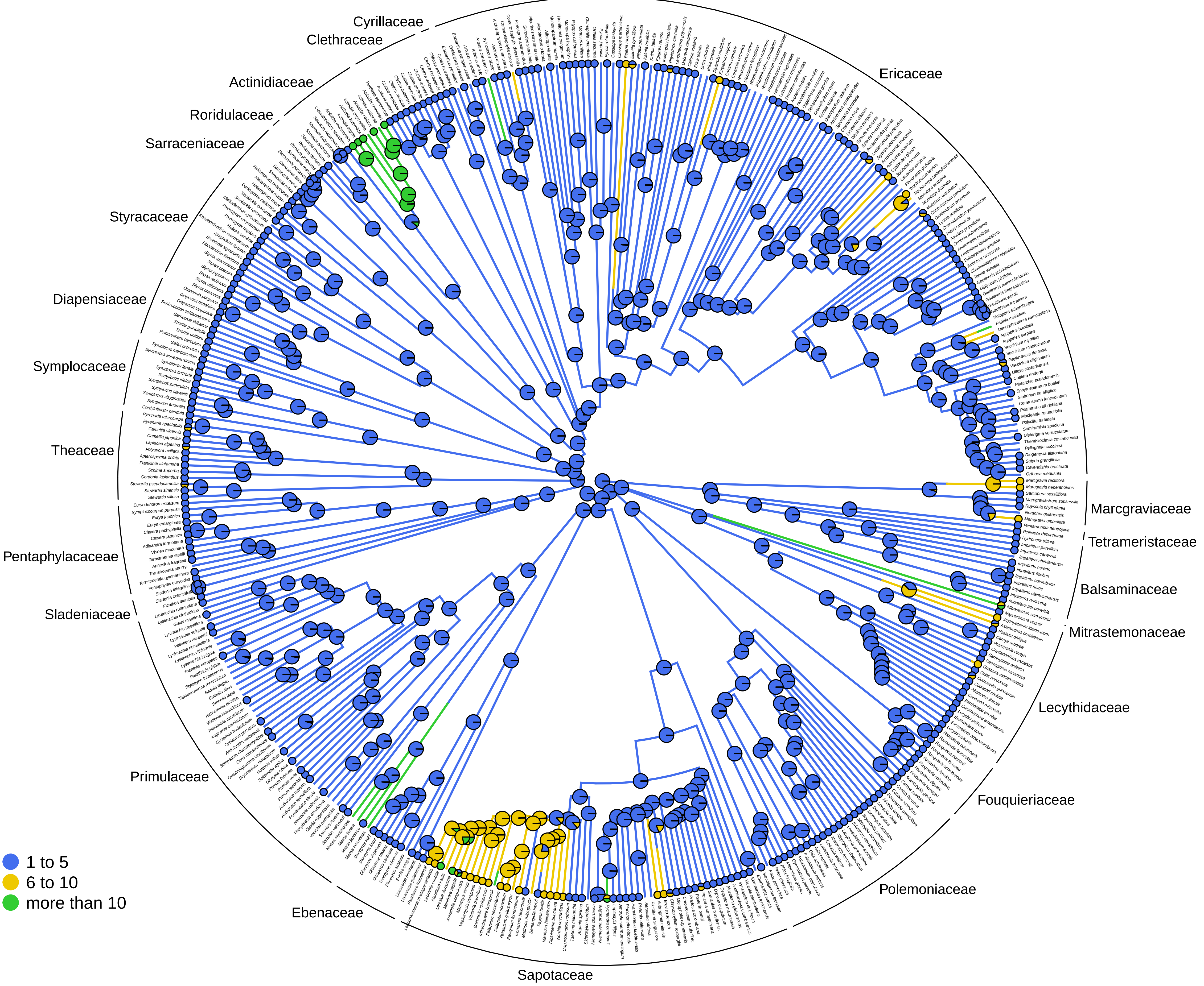

#### Number of fertile stamens

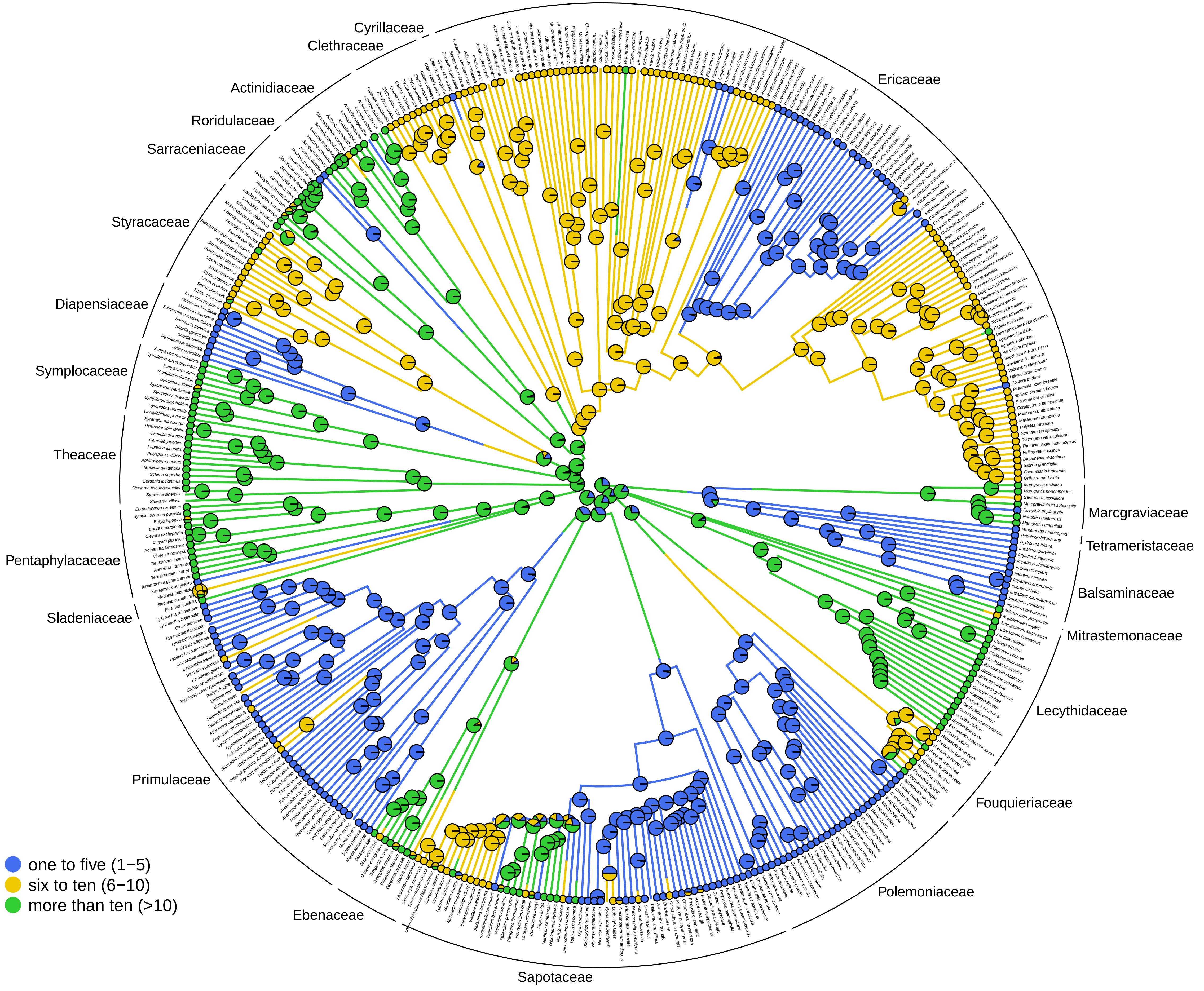

### Number of ovules per functional carpel

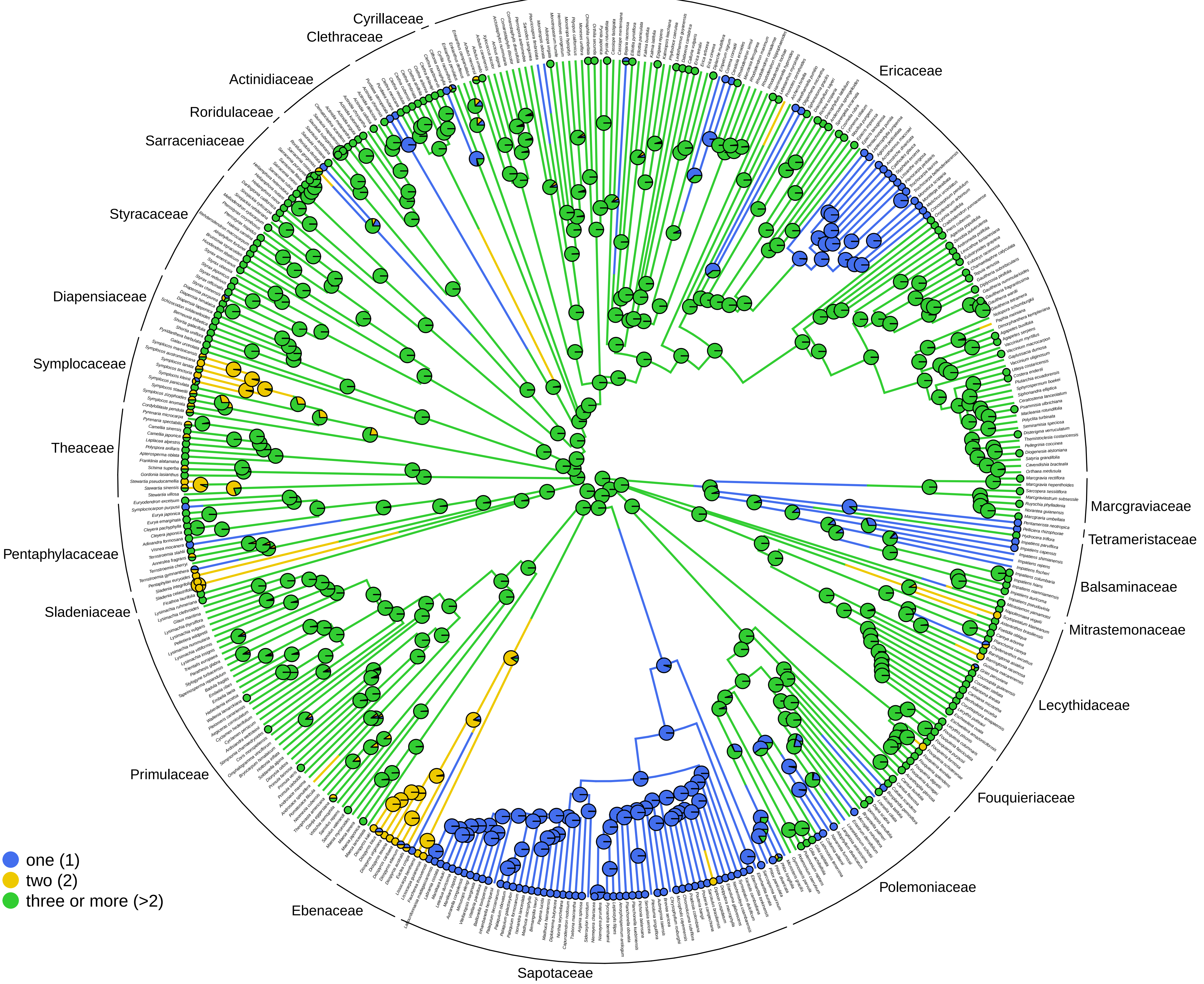

### Number of Petals

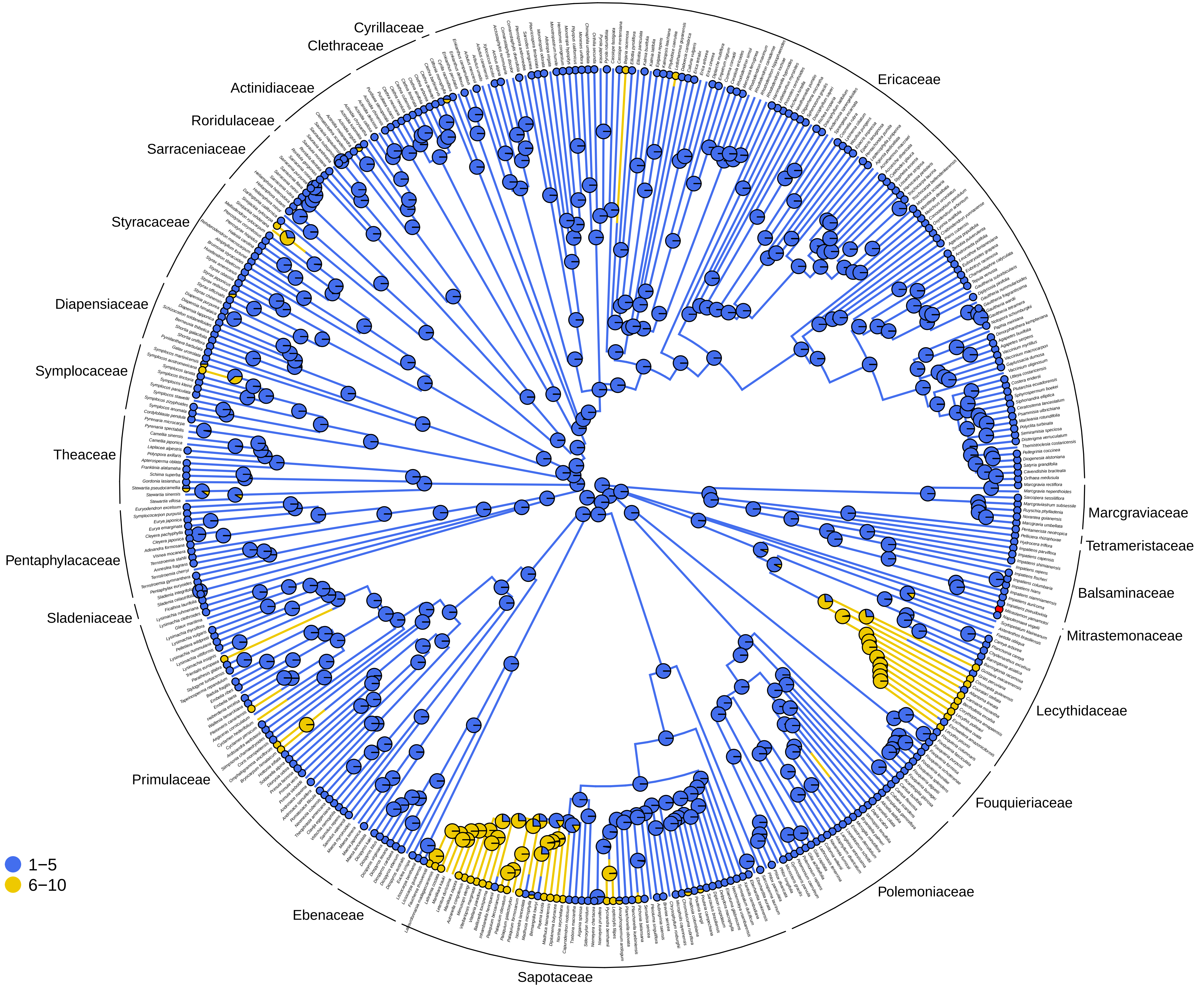

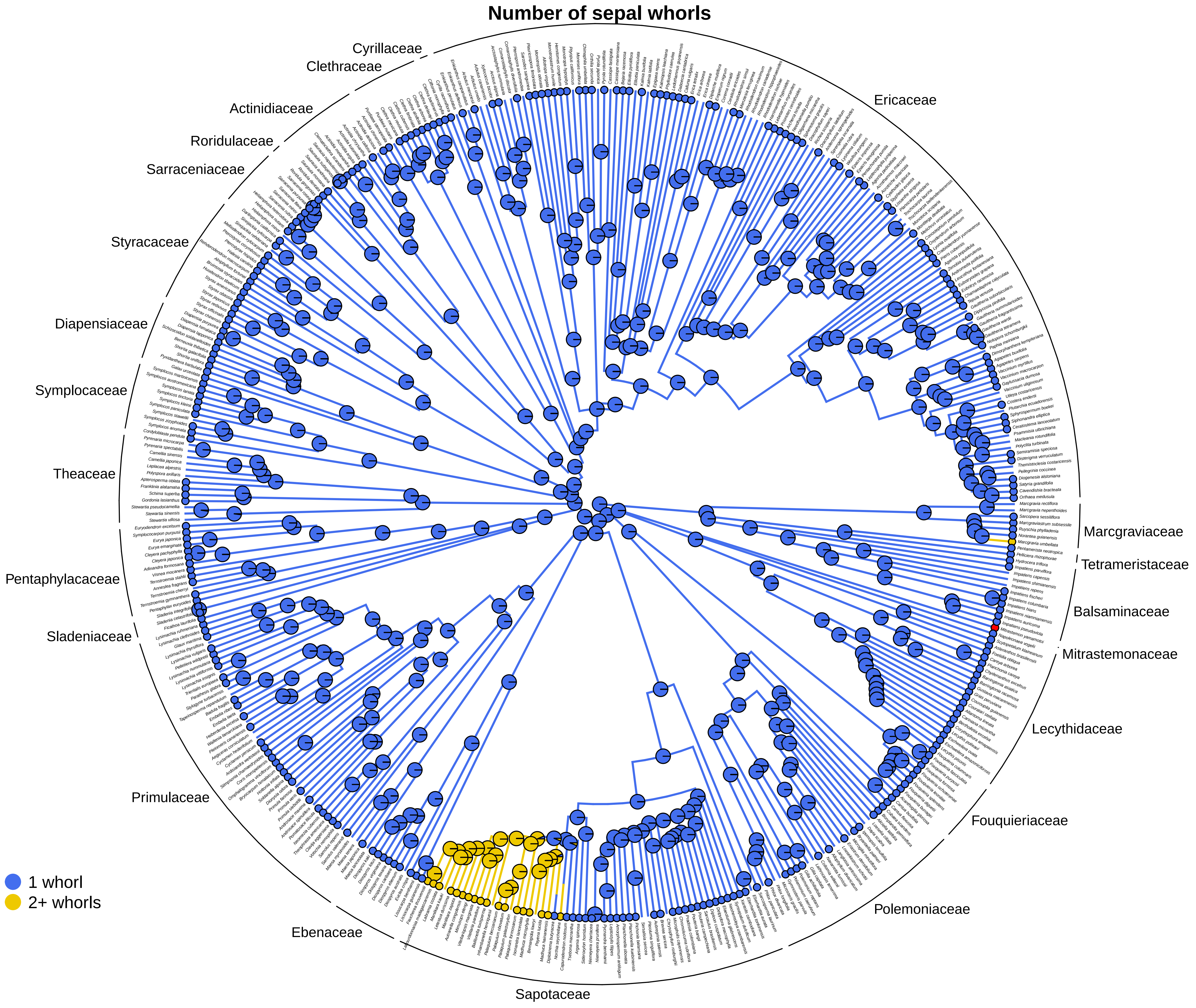
Number of sepal whorls
Cyrillaceae
Clethraceae
Ericaceae
Actinidiaceae
Roridulaceae
Sarraceniaceae
Styracaceae
Diapensiaceae
Symplocaceae
Theaceae
Pentaphylacaceae
Sladeniaceae
Primulaceae
Ebenaceae
Sapotaceae
Polemoniaceae
Fouquieriaceae
Lecythidaceae
Mitrastemonaceae
Balsaminaceae
Tetrameristaceae
Marcgraviaceae
1 whorl
2+ whorls

### Number of Sepals

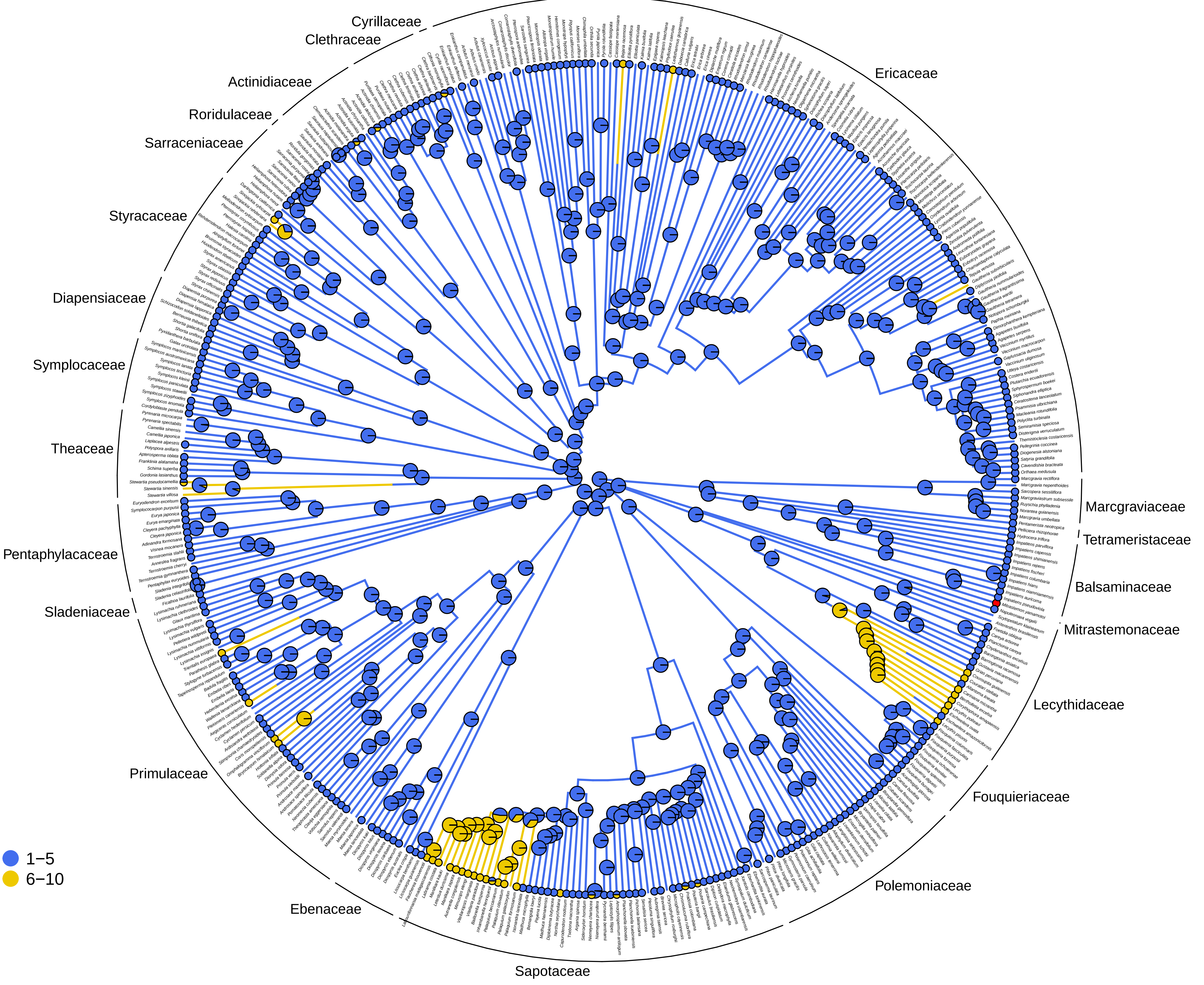

#### Ovary position

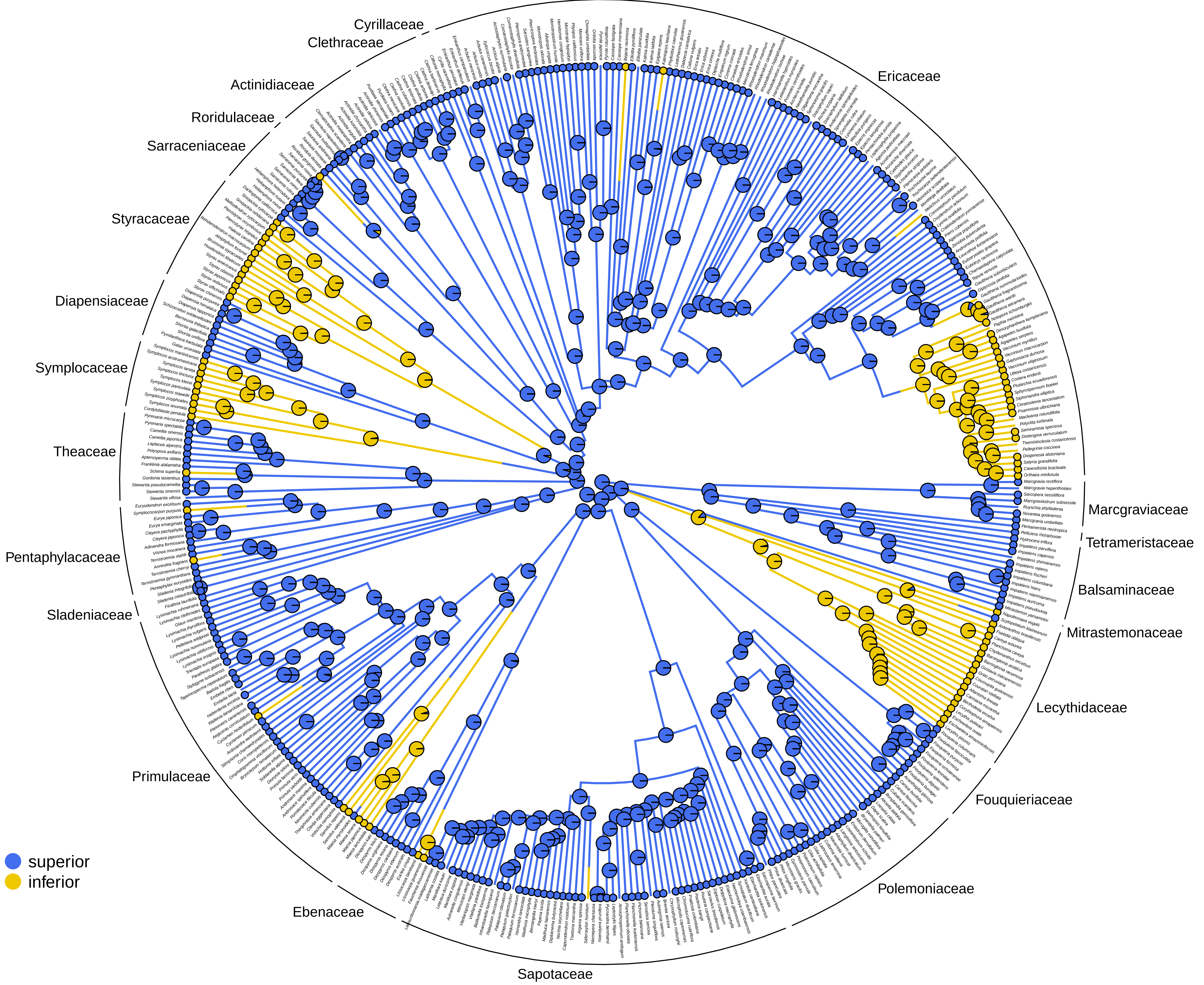

### Ovule integument

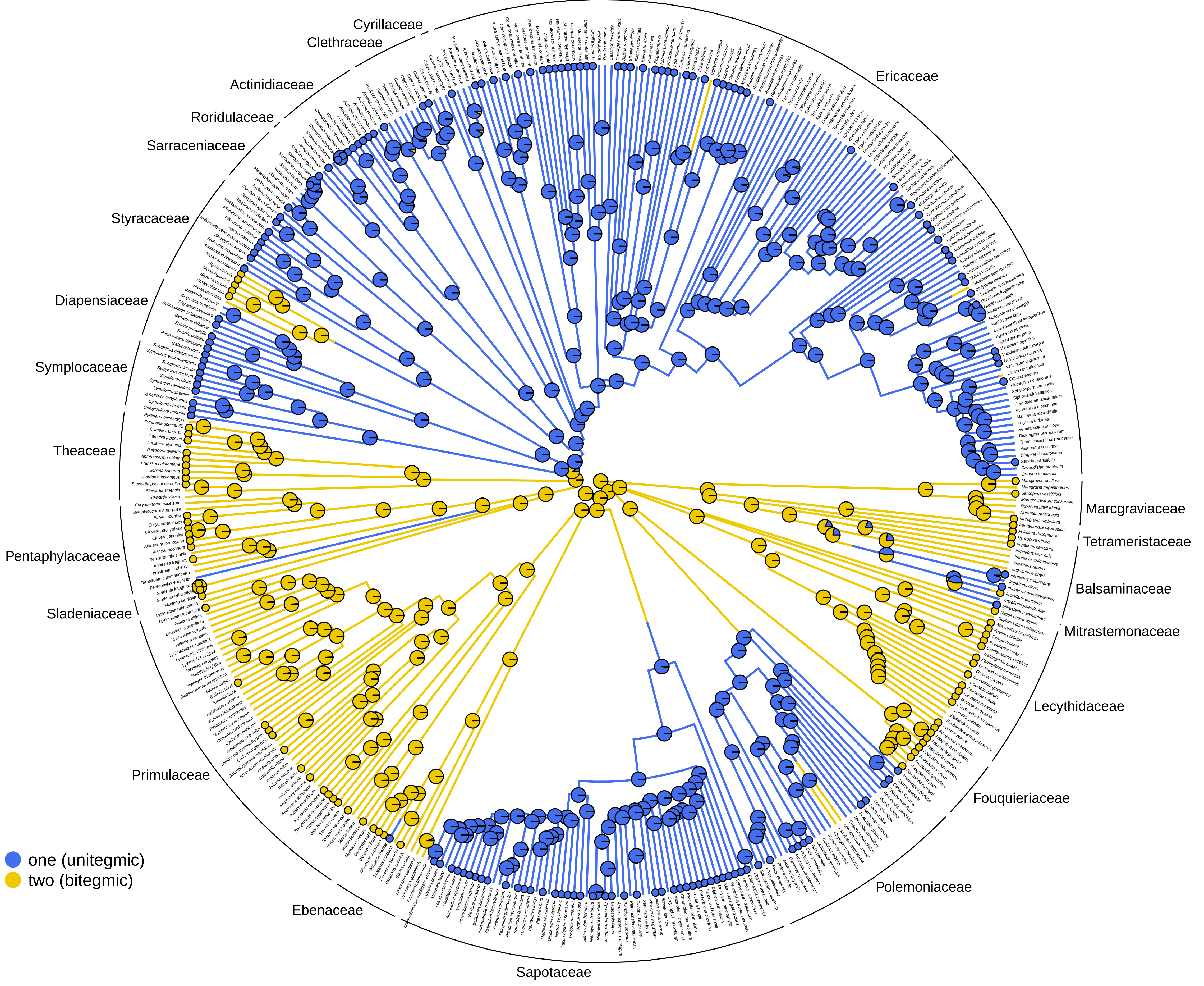

**Perianth differentiation**

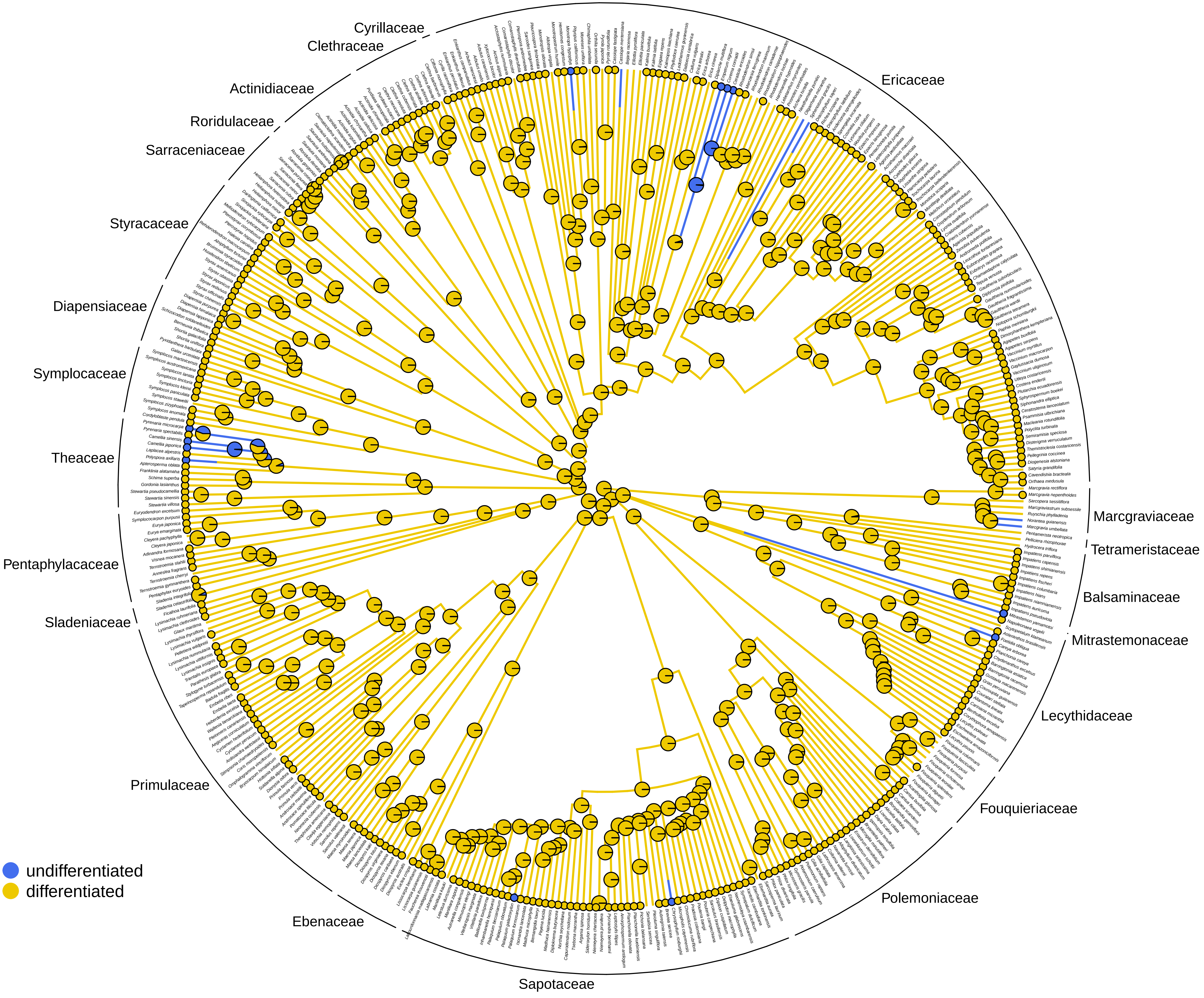

**Petal aestivation**

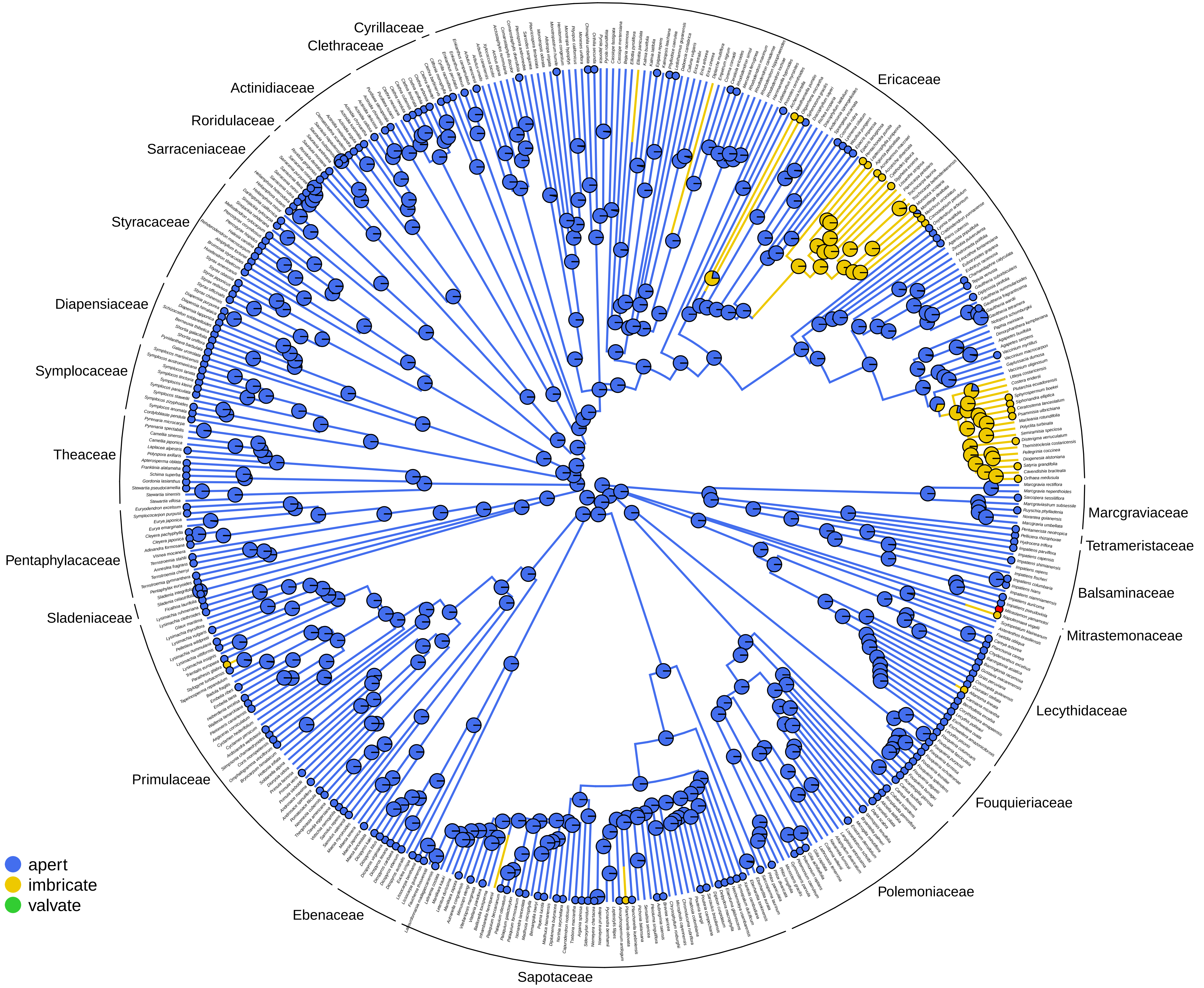

### Petal phyllotaxis

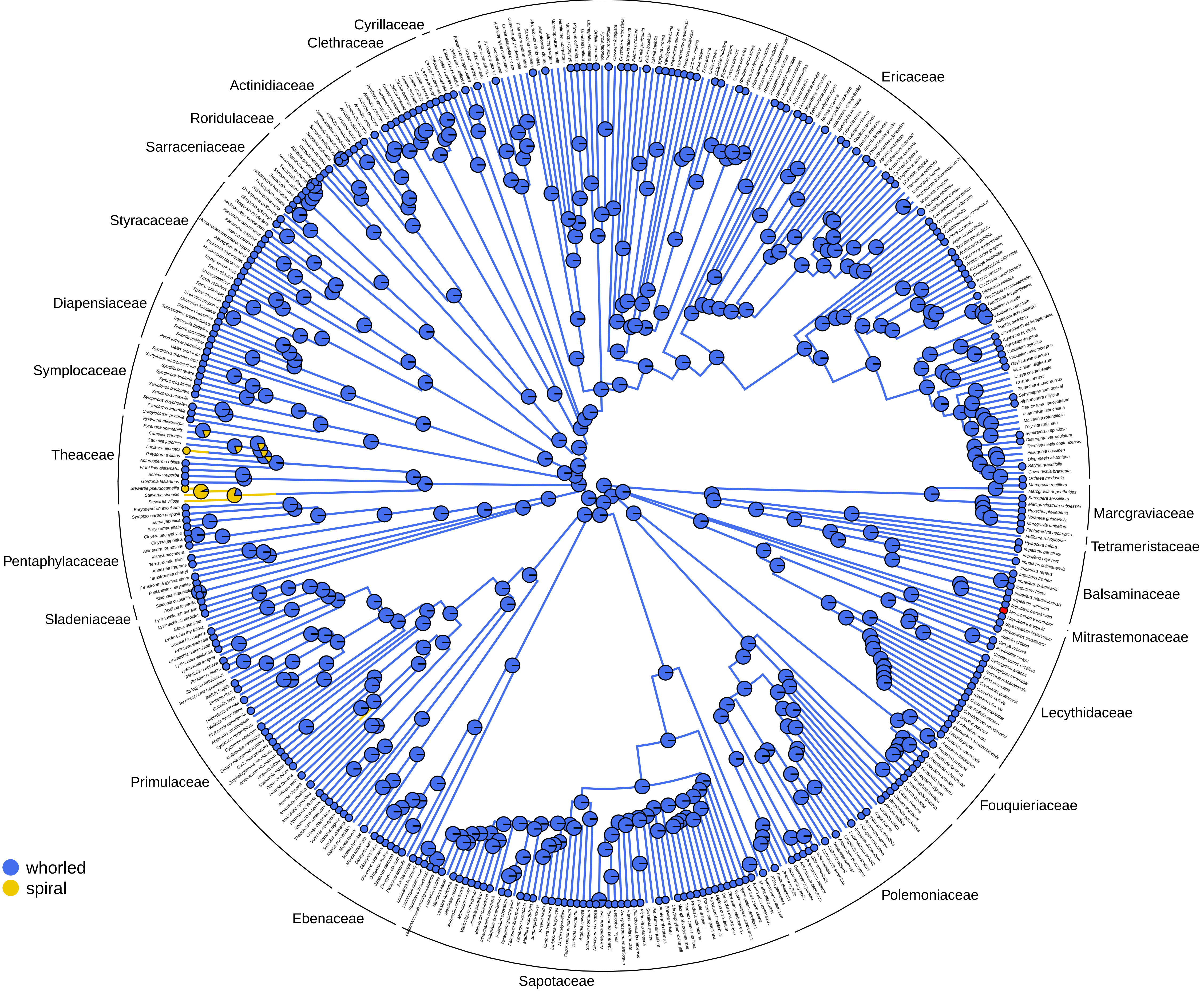

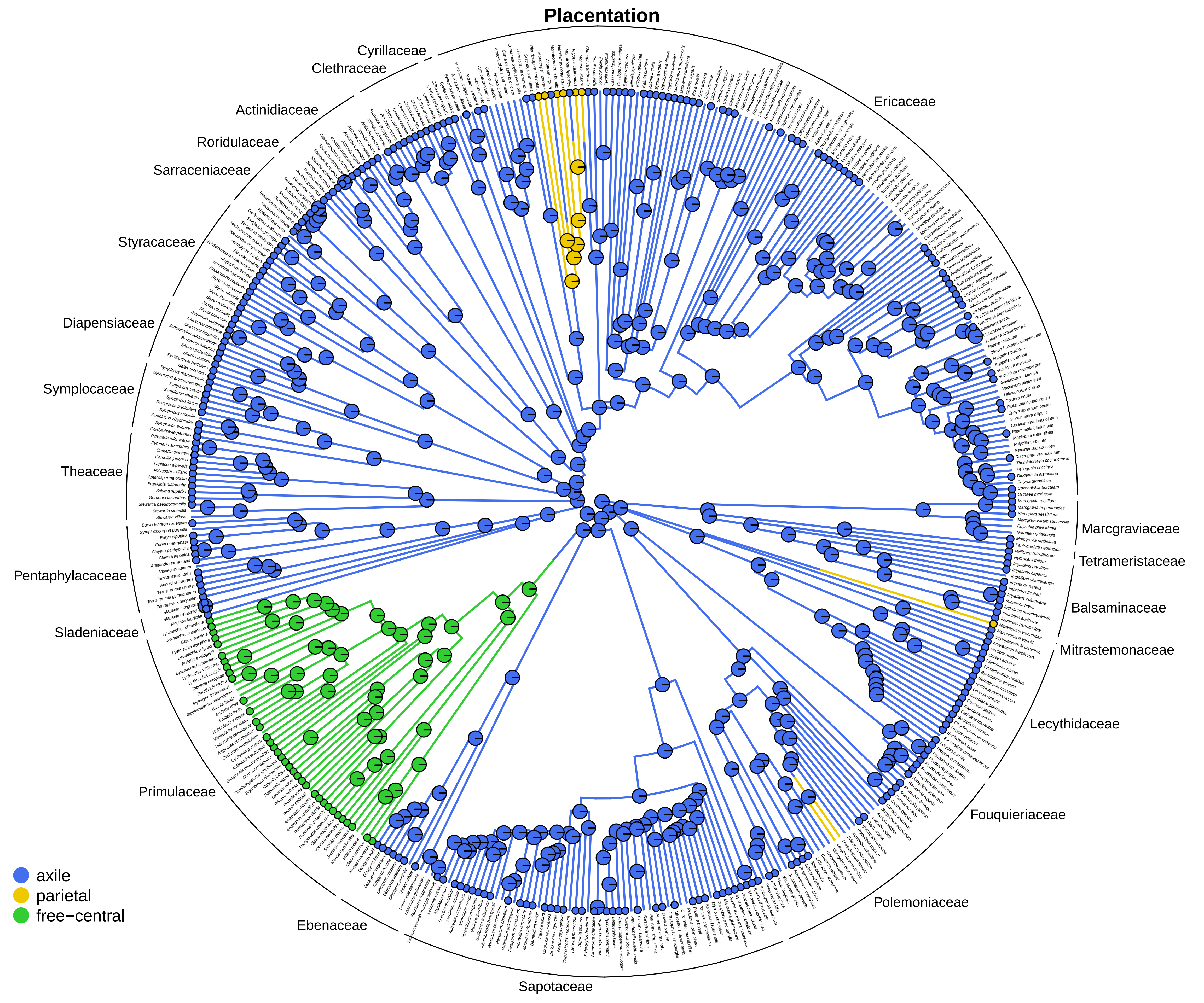
Placentation
Cyrillaceae
Clethraceae
Actinidiaceae
Roridulaceae
Sarraceniaceae
Styracaceae
Diapensiaceae
Symplocaceae
Theaceae
Pentaphylacaceae
Sladeniaceae
Primulaceae
Ebenaceae
Sapotaceae
Polemoniaceae
Fouquieriaceae
Lecythidaceae
Mitrastemonaceae
Balsaminaceae
Tetrameristaceae
Marcgraviaceae
Ericaceae
axile
parietal
free−central

#### Presence of staminodes

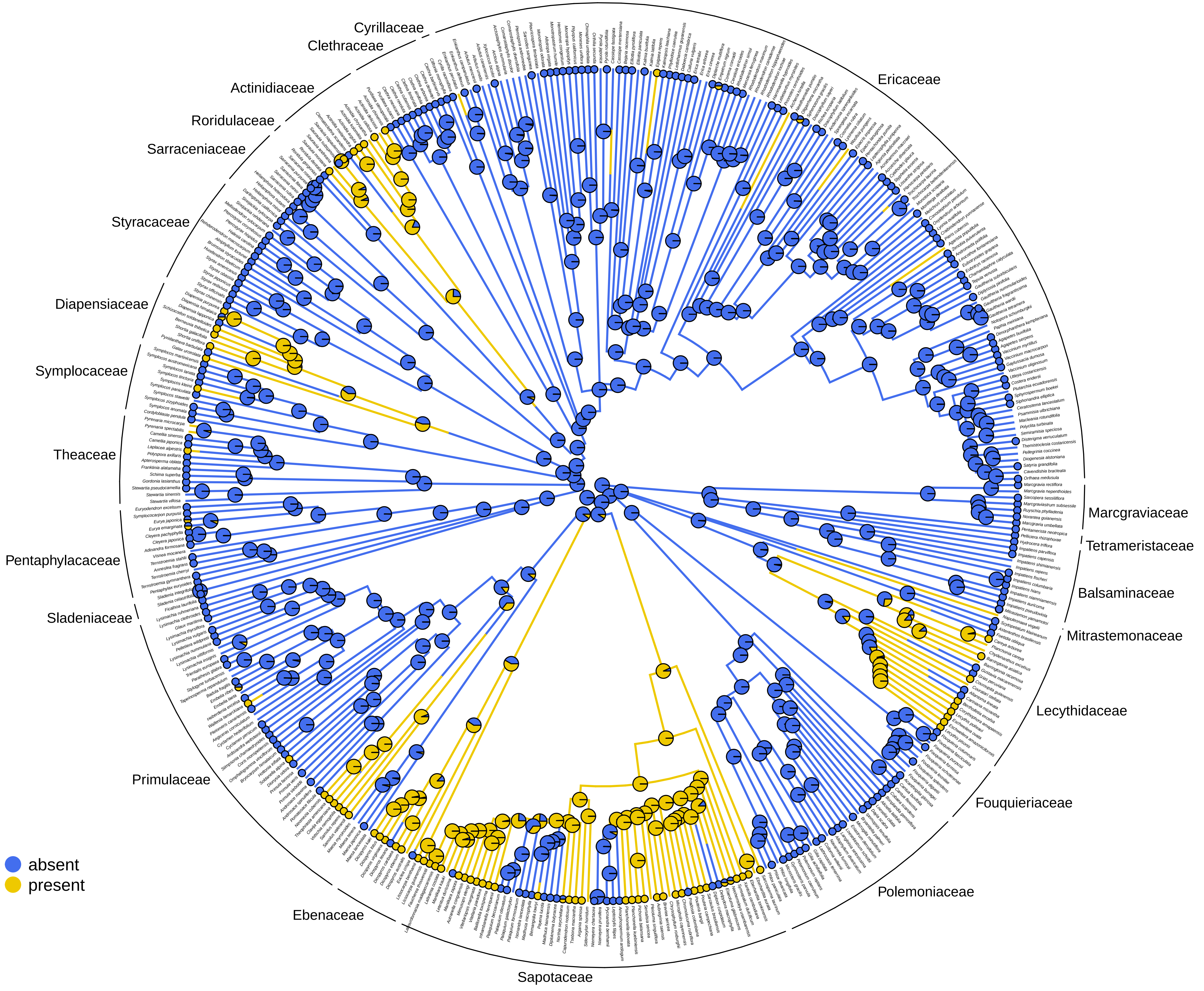

### Sepal aestivation

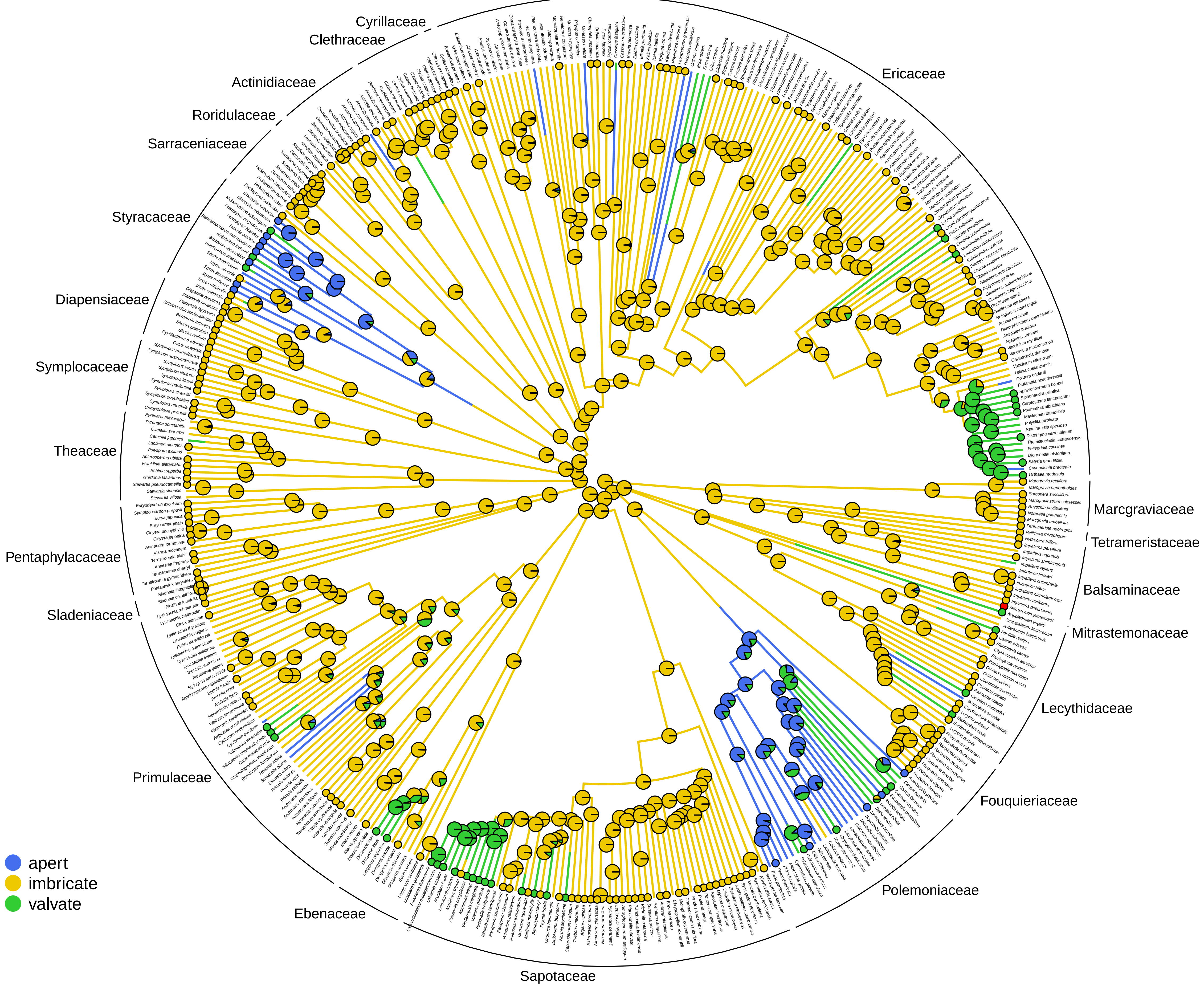

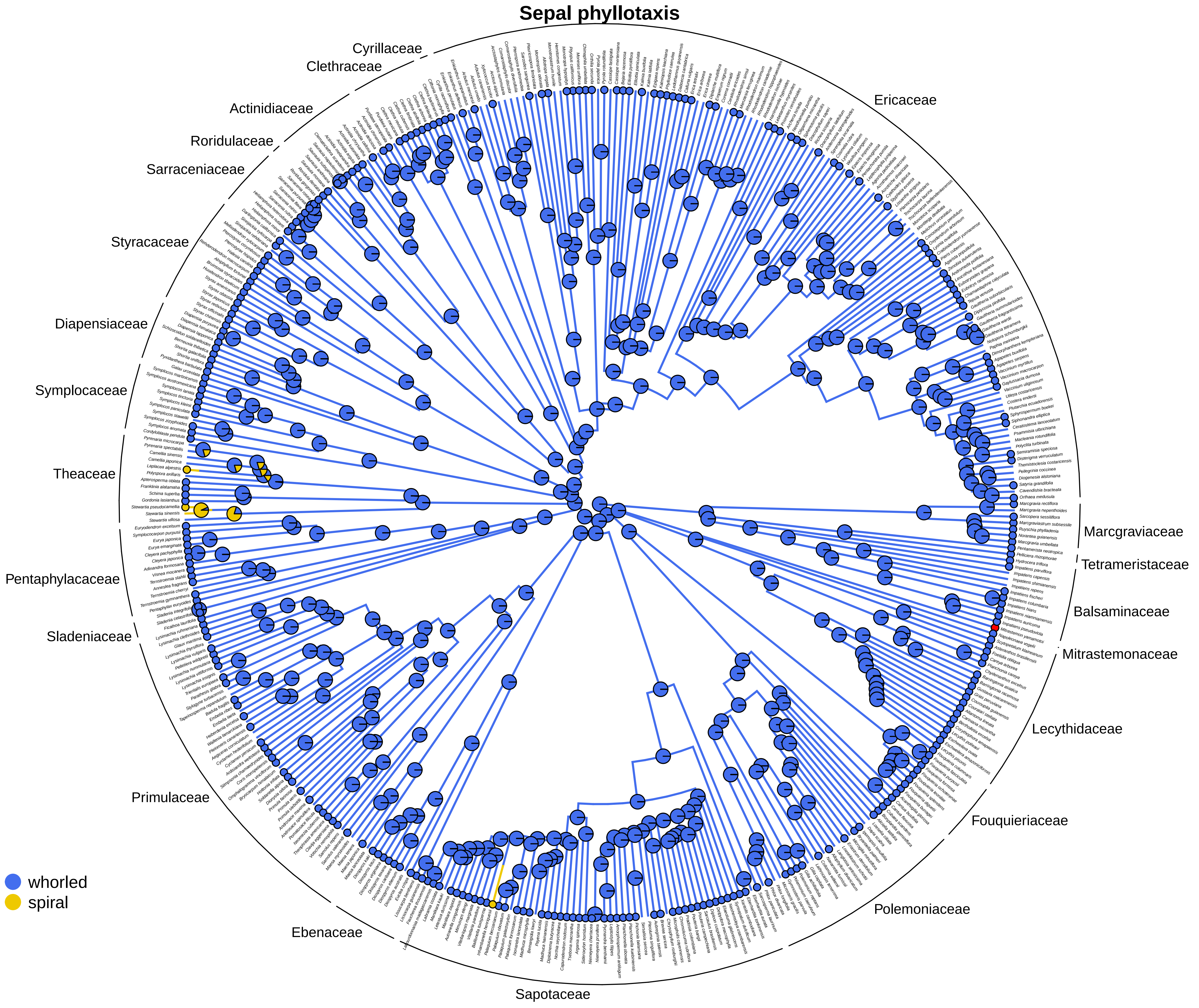
Sepal phyllotaxis
Cyrillaceae
Clethraceae
Ericaceae
Actinidiaceae
Roridulaceae
Sarraceniaceae
Styracaceae
Diapensiaceae
Symplocaceae
Theaceae
Pentaphylacaceae
Sladeniaceae
Primulaceae
Ebenaceae
Sapotaceae
Polemoniaceae
Fouquieriaceae
Lecythidaceae
Mitrastemonaceae
Balsaminaceae
Tetrameristaceae
Marcgraviaceae
whorled
spiral

#### Structural sex of flowers

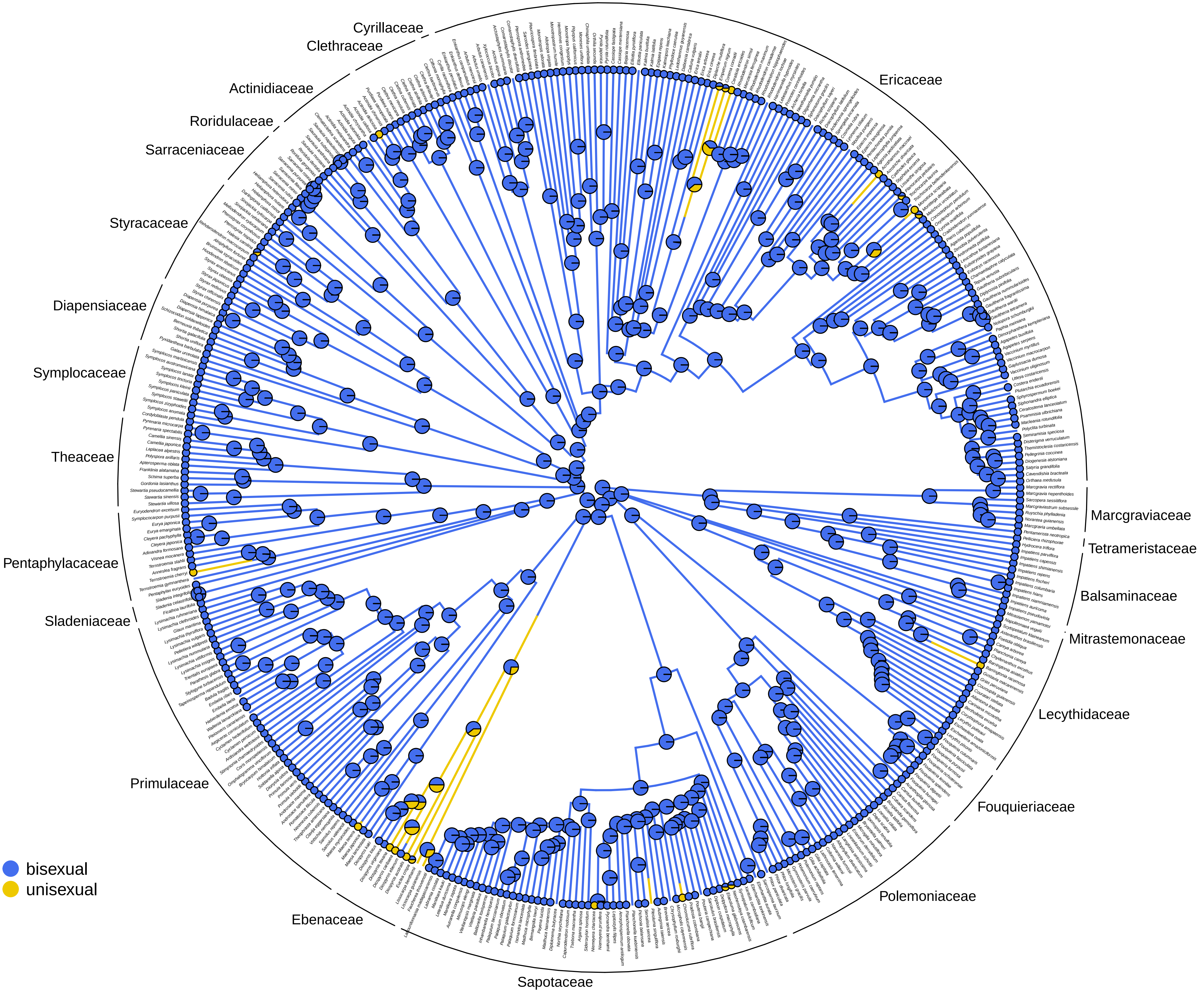

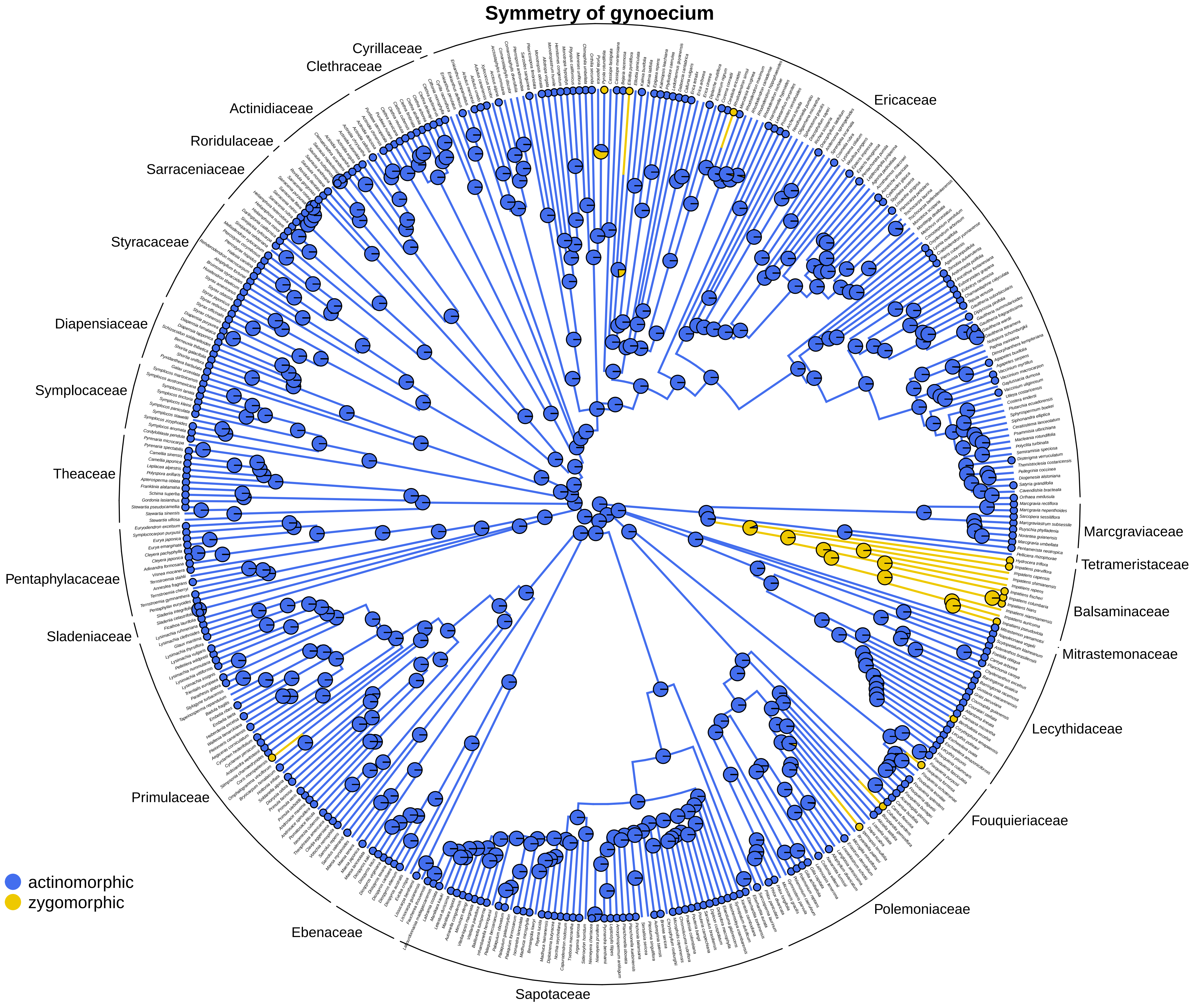
Symmetry of gynoecium
Cyrillaceae
Clethraceae
Ericaceae
Actinidiaceae
Roridulaceae
Sarraceniaceae
Styracaceae
Diapensiaceae
Symplocaceae
Theaceae
Pentaphylacaceae
Sladeniaceae
Primulaceae
Ebenaceae
Sapotaceae
Polemoniaceae
Fouquieriaceae
Lecythidaceae
Mitrastemonaceae
Balsaminaceae
Tetrameristaceae
Marcgraviaceae
actinomorphic
zygomorphic

#### Symmetry of perianth

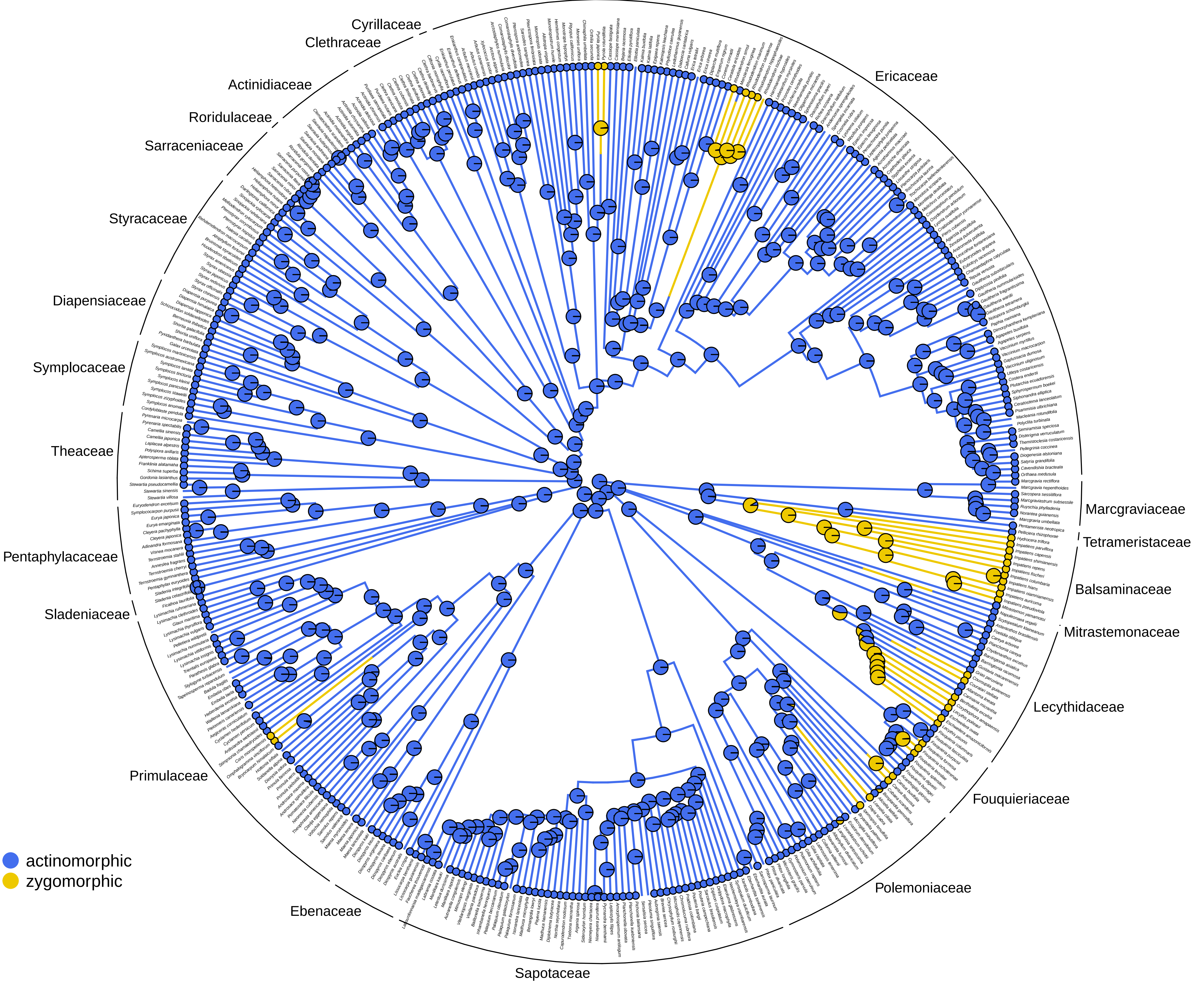

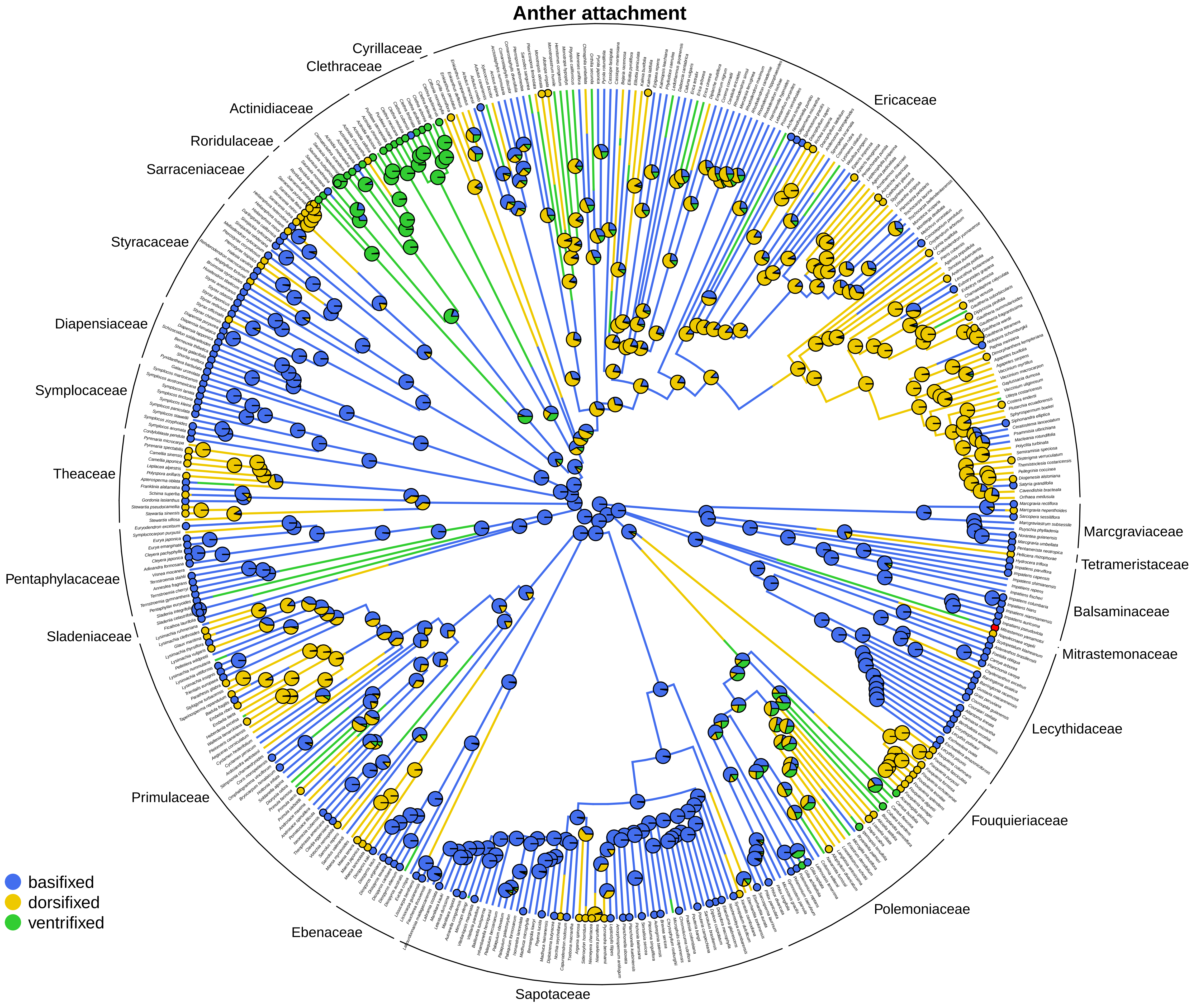
Anther attachment
Cyrillaceae
Clethraceae
Ericaceae
Actinidiaceae
Roridulaceae
Sarraceniaceae
Styracaceae
Diapensiaceae
Symplocaceae
Theaceae
Pentaphylacaceae
Sladeniaceae
Primulaceae
Ebenaceae
Sapotaceae
Polemoniaceae
Fouquieriaceae
Lecythidaceae
Mitrastemonaceae
Balsaminaceae
Tetrameristaceae
Marcgraviaceae
basifixed
dorsifixed
ventrifixed

**Anther orientation**

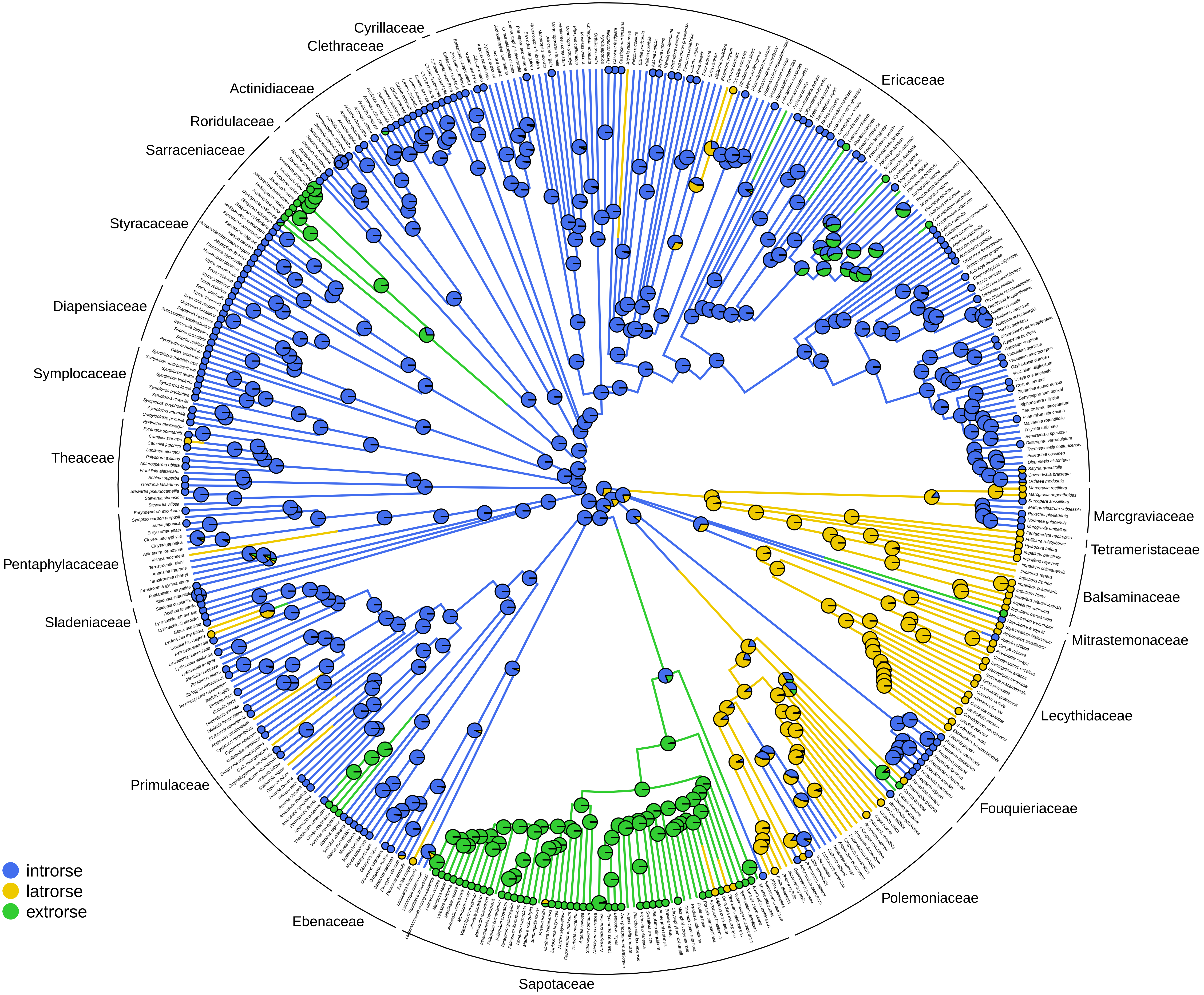

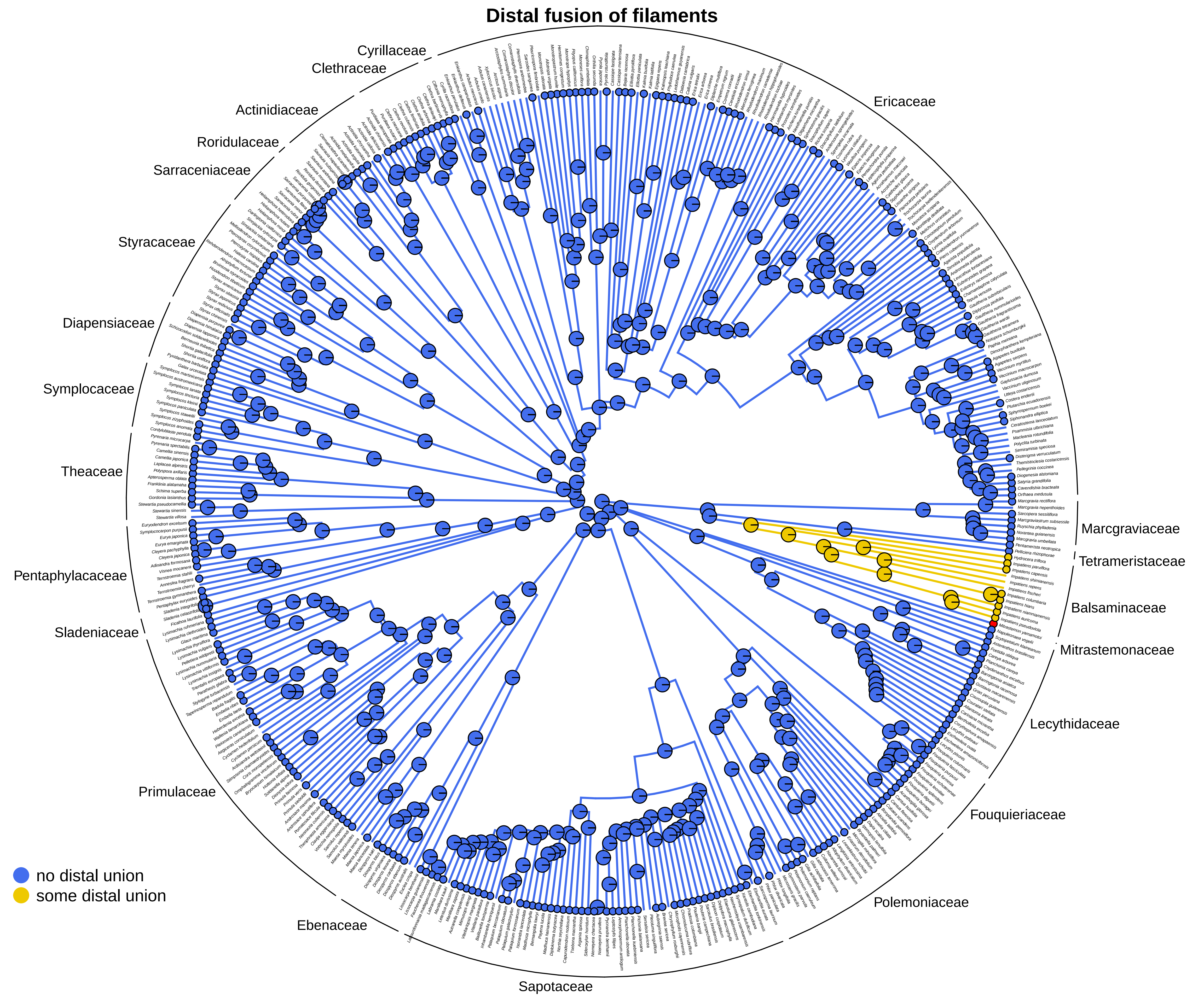
Distal fusion of filaments
Cyrillaceae
Clethraceae
Actinidiaceae
Roridulaceae
Sarraceniaceae
Styracaceae
Diapensiaceae
Symplocaceae
Theaceae
Pentaphylacaceae
Sladeniaceae
Primulaceae
Ebenaceae
Sapotaceae
Polemoniaceae
Fouquieriaceae
Lecythidaceae
Mitrastemonaceae
Balsaminaceae
Tetrameristaceae
Marcgraviaceae
Ericaceae
no distal union
some distal union

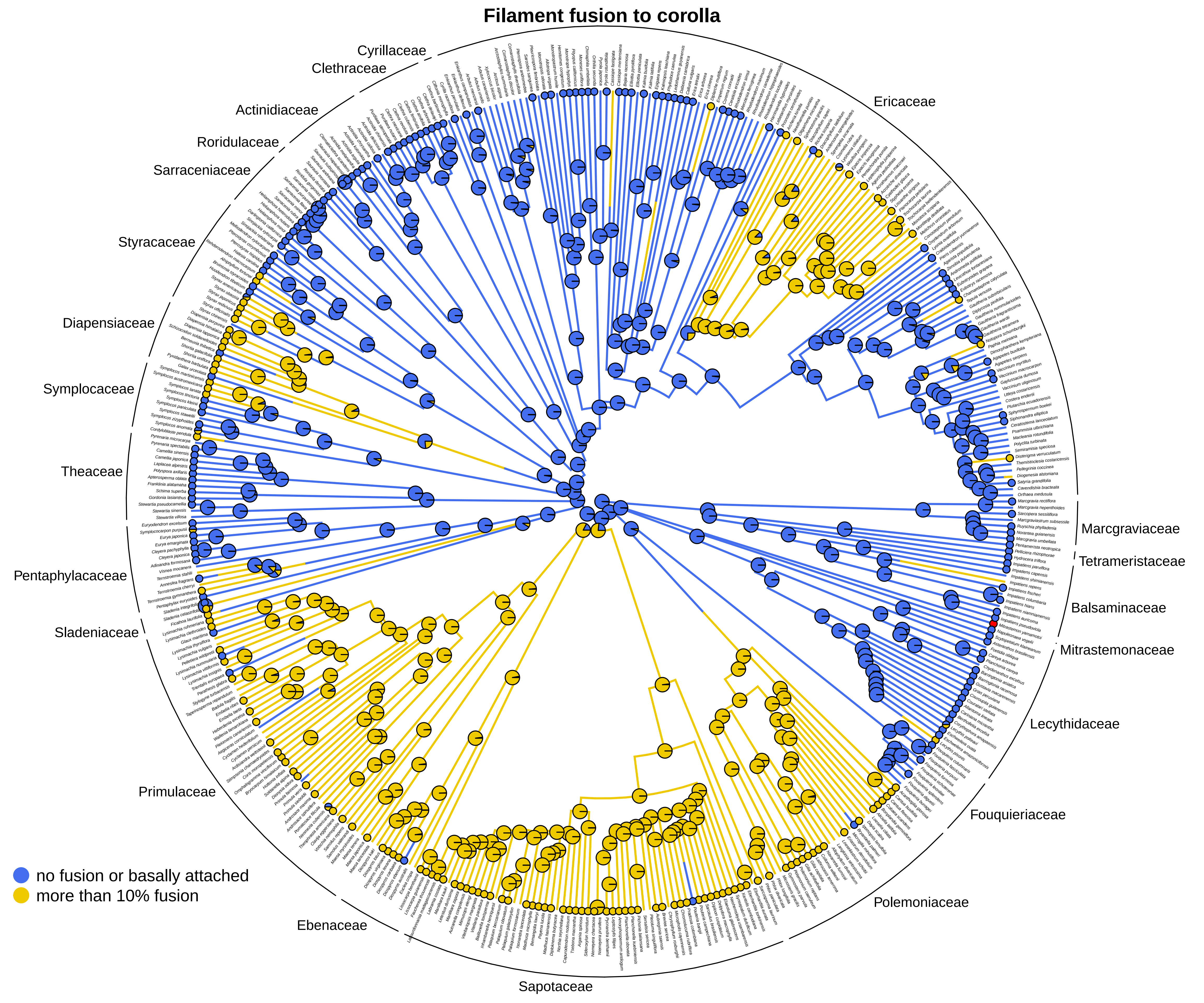
Filament fusion to corolla
Cyrillaceae
Clethraceae
Actinidiaceae
Roridulaceae
Sarraceniaceae
Styracaceae
Diapensiaceae
Symplocaceae
Theaceae
Pentaphylacaceae
Sladeniaceae
Primulaceae
Ebenaceae
Sapotaceae
Polemoniaceae
Fouquieriaceae
Lecythidaceae
Mitrastemonaceae
Balsaminaceae
Tetrameristaceae
Marcgraviaceae
Ericaceae
no fusion or basally attached
more than 10% fusion

### Filament insertion to corolla

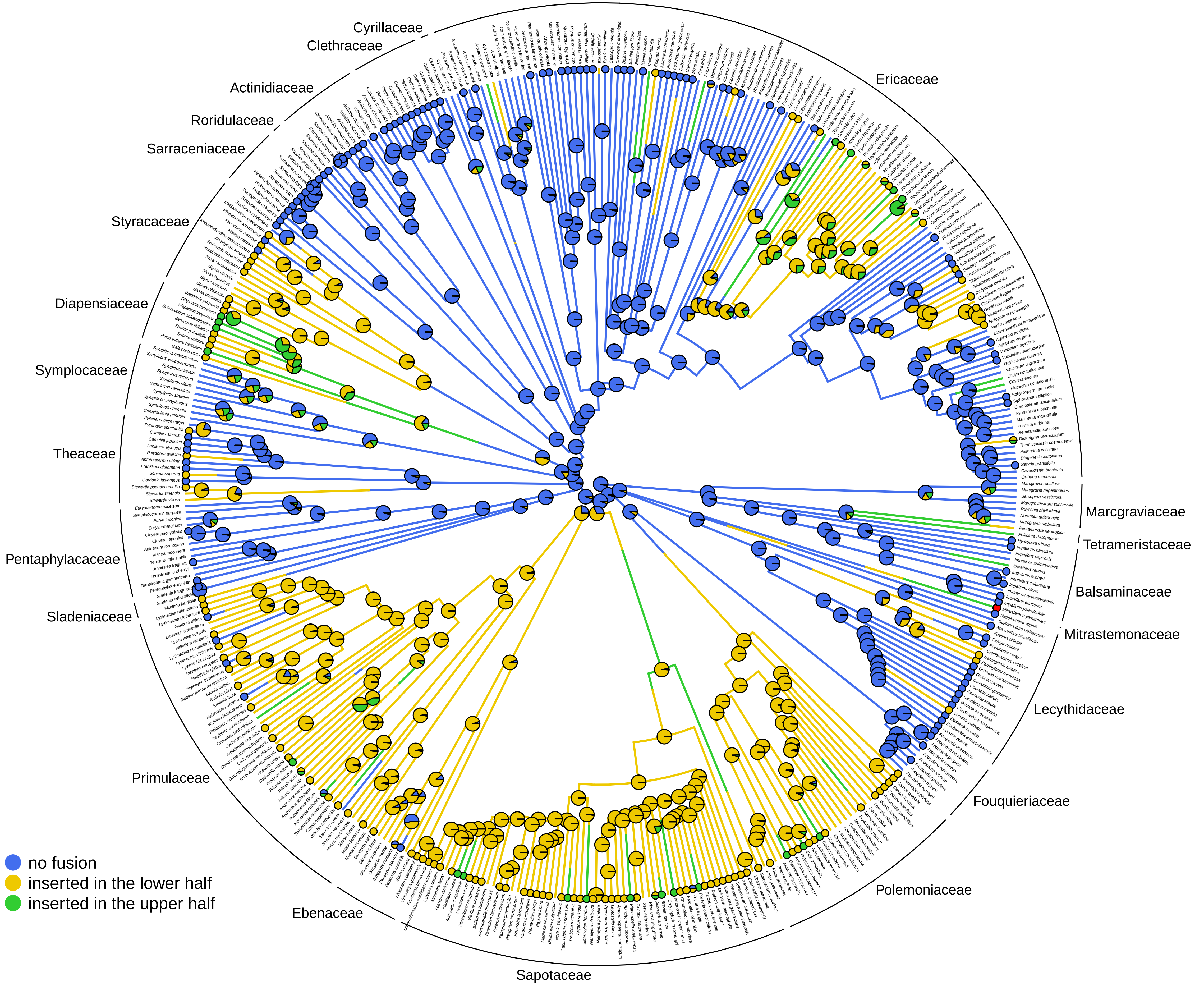

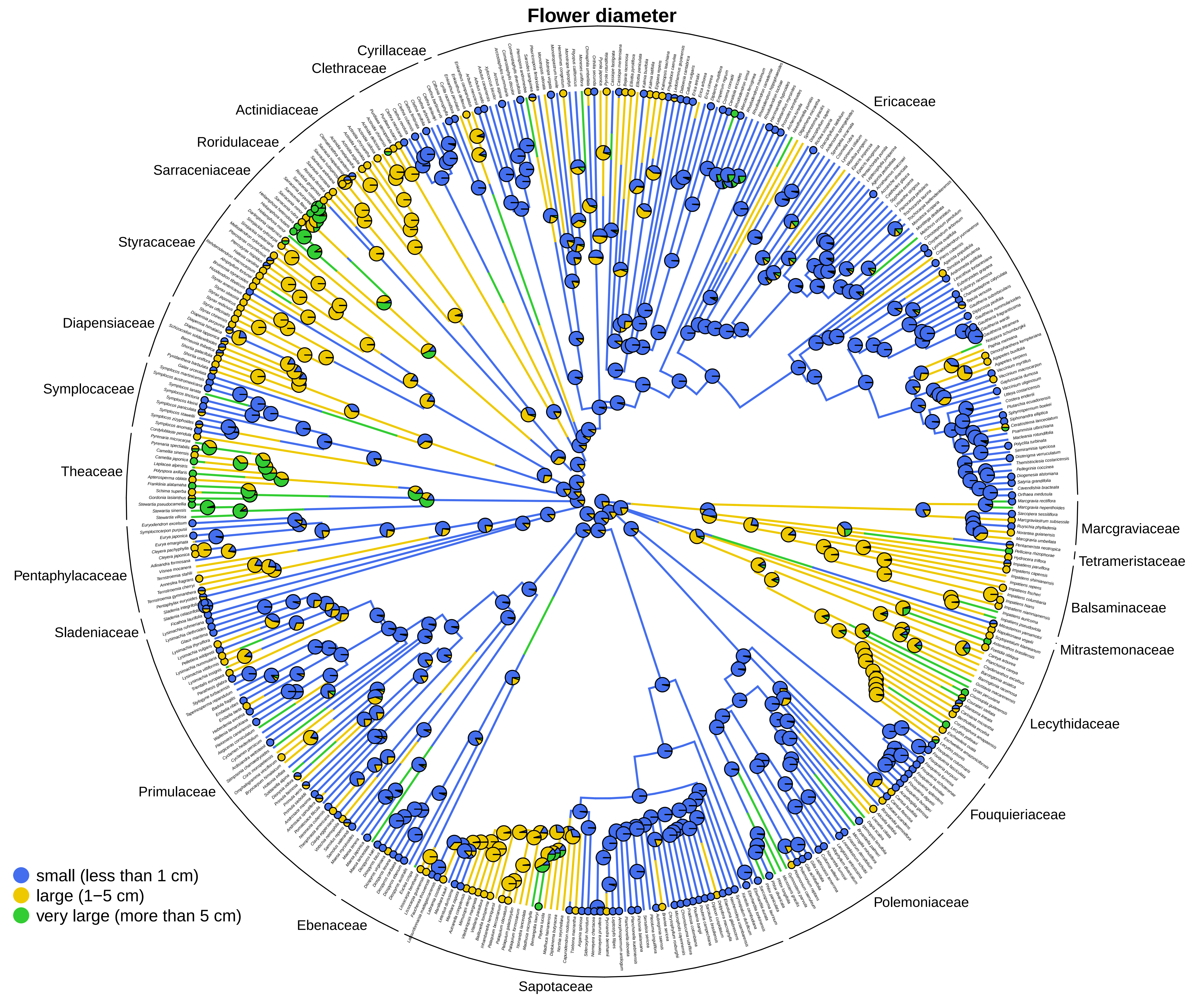
Flower diameter
Cyrillaceae
Clethraceae
Ericaceae
Actinidiaceae
Roridulaceae
Sarraceniaceae
Styracaceae
Diapensiaceae
Symplocaceae
Theaceae
Pentaphylacaceae
Sladeniaceae
Primulaceae
Ebenaceae
Sapotaceae
Polemoniaceae
Fouquieriaceae
Lecythidaceae
Mitrastemonaceae
Balsaminaceae
Tetrameristaceae
Marcgraviaceae
small (less than 1 cm)
large (1−5 cm)
very large (more than 5 cm)

### Flower length

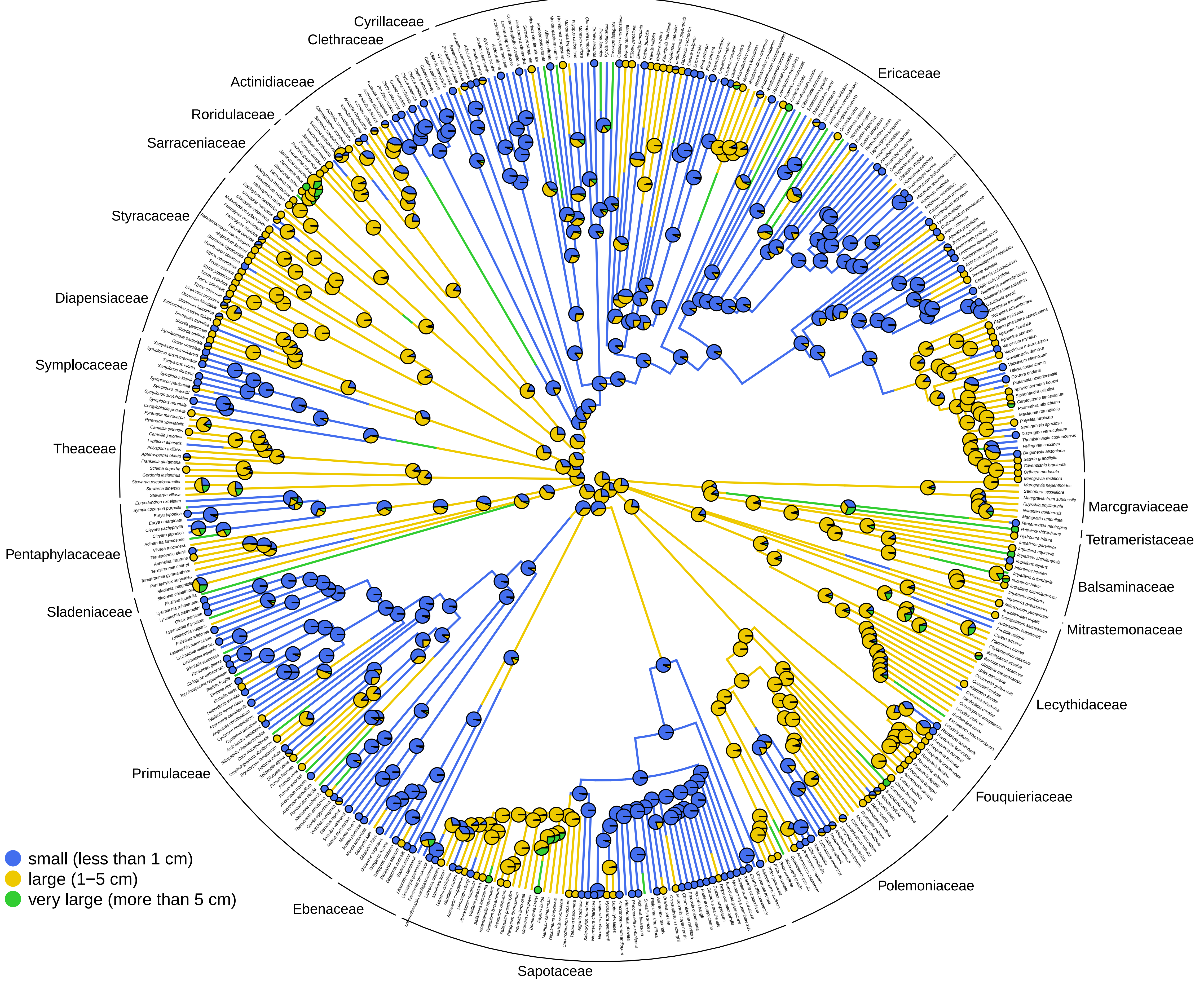

### Fusion of anthers

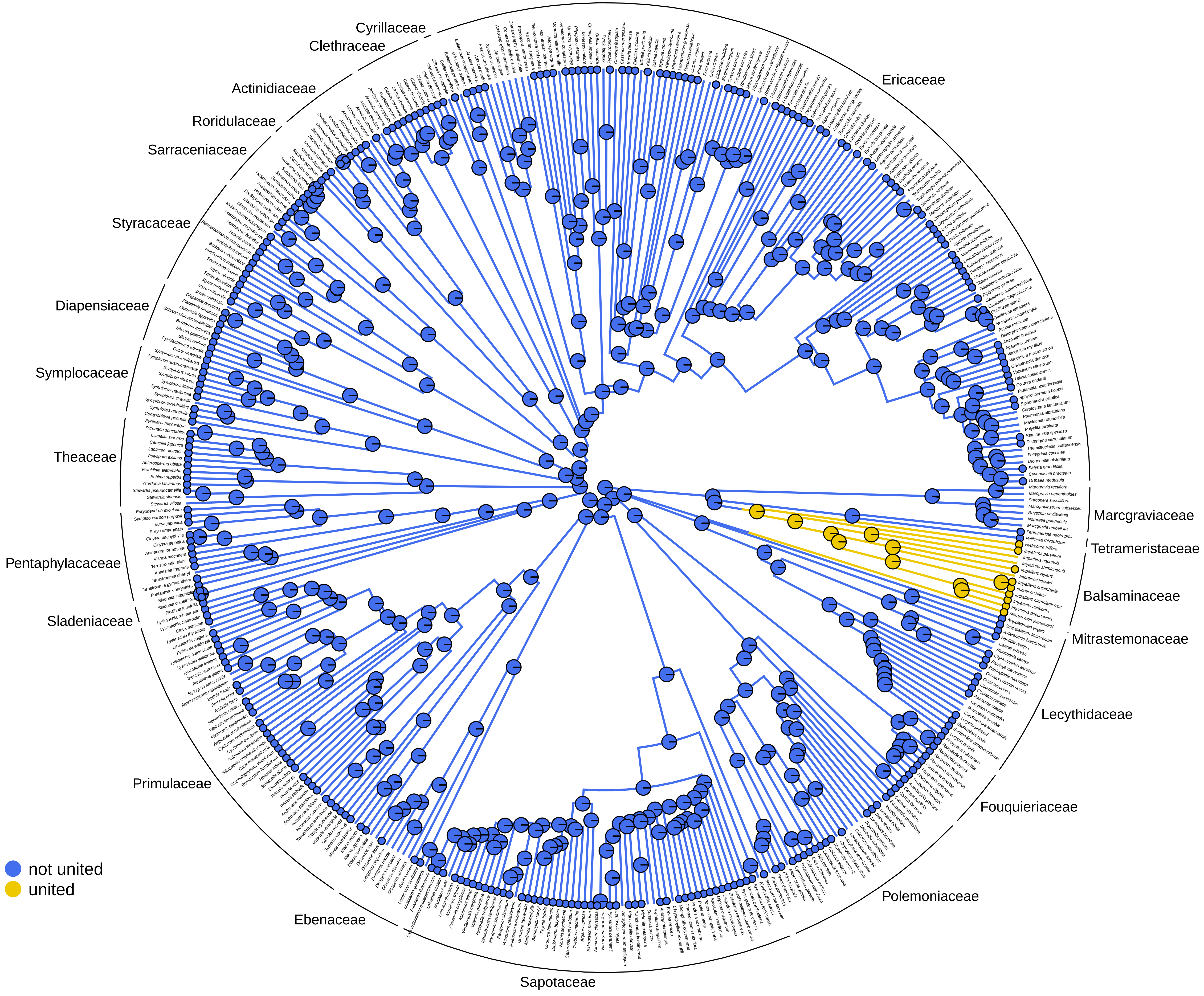

### Fusion of filaments

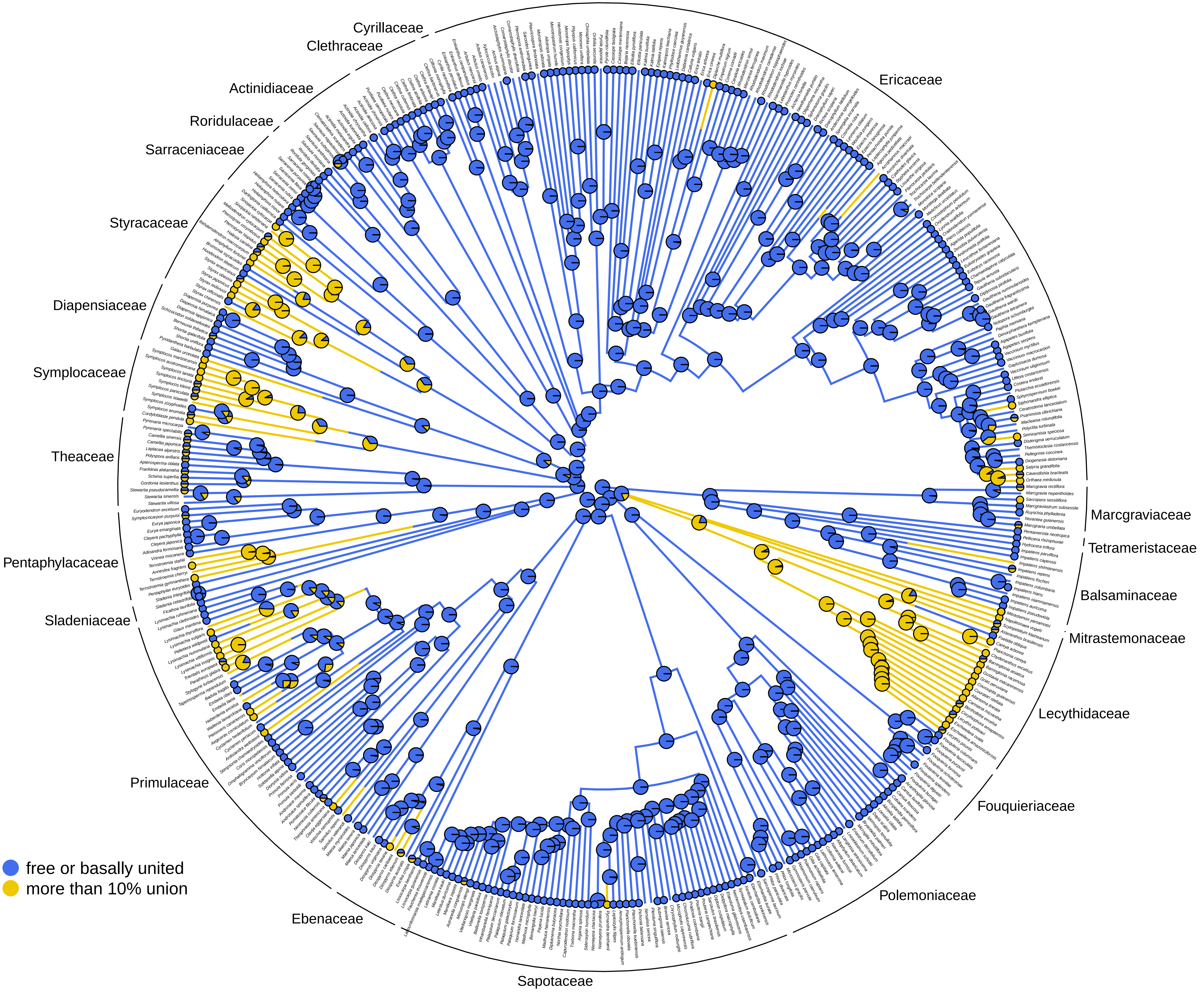

### Fusion of petals

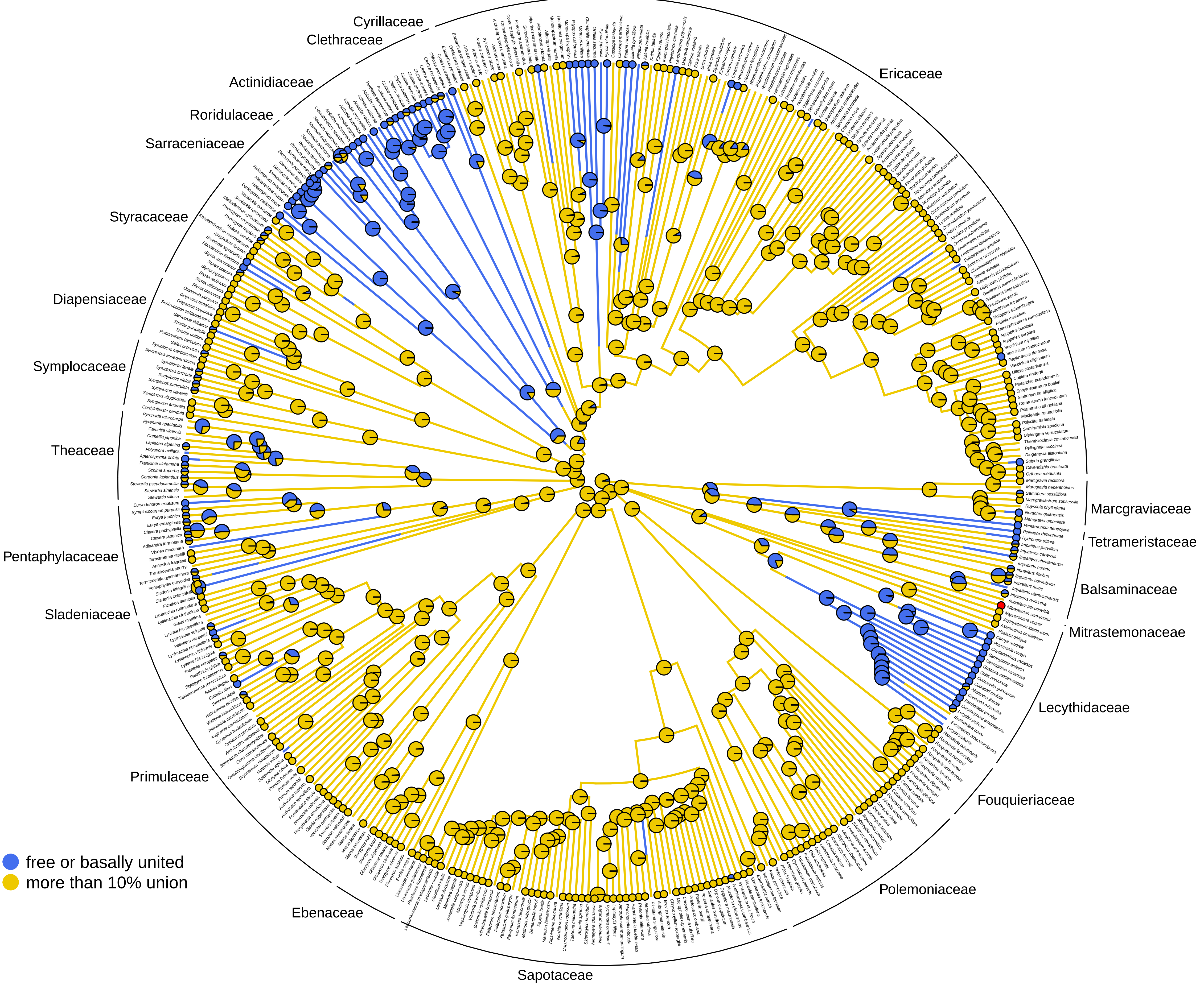

### Fusion of sepals

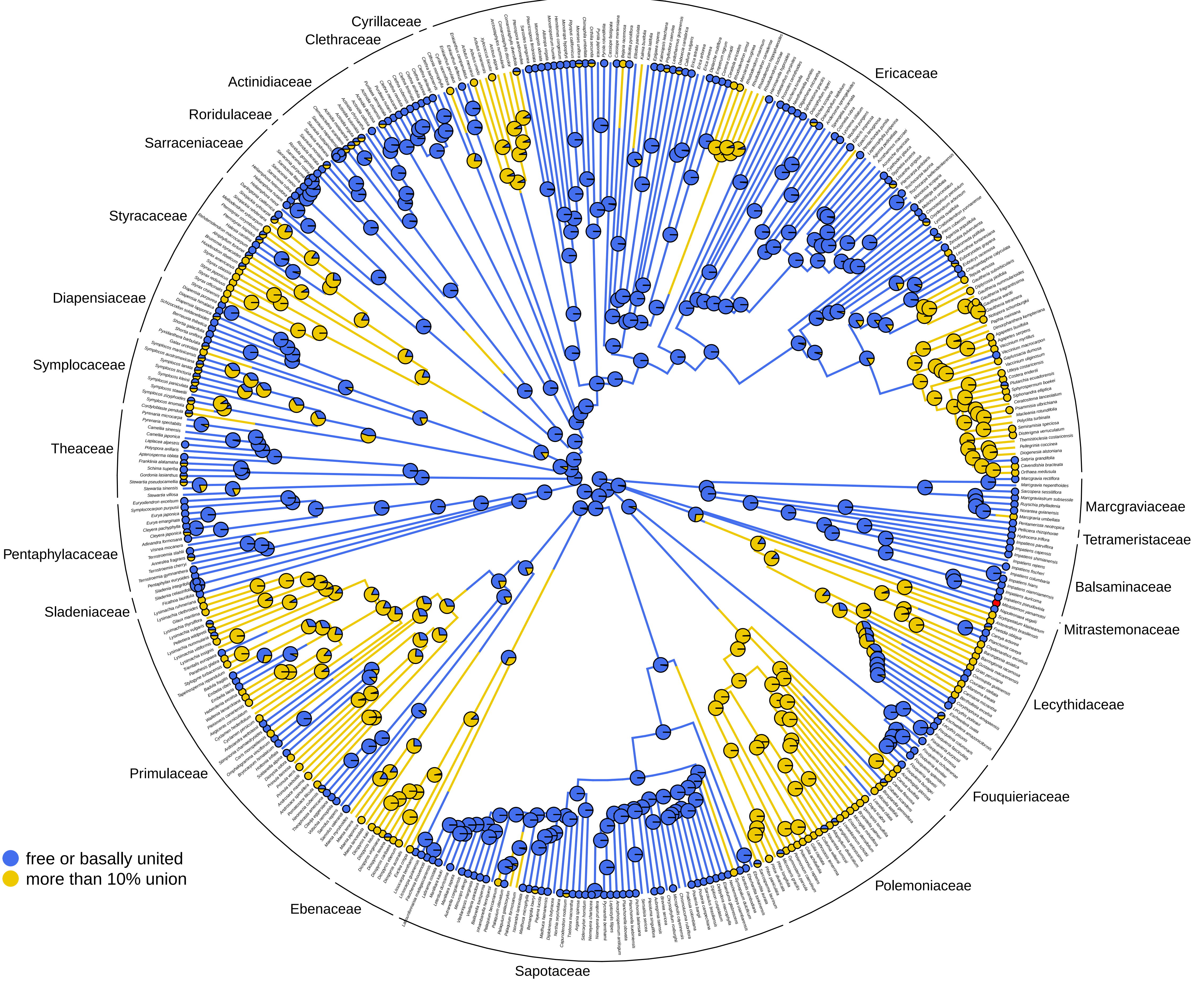

### Fusion of stigmas

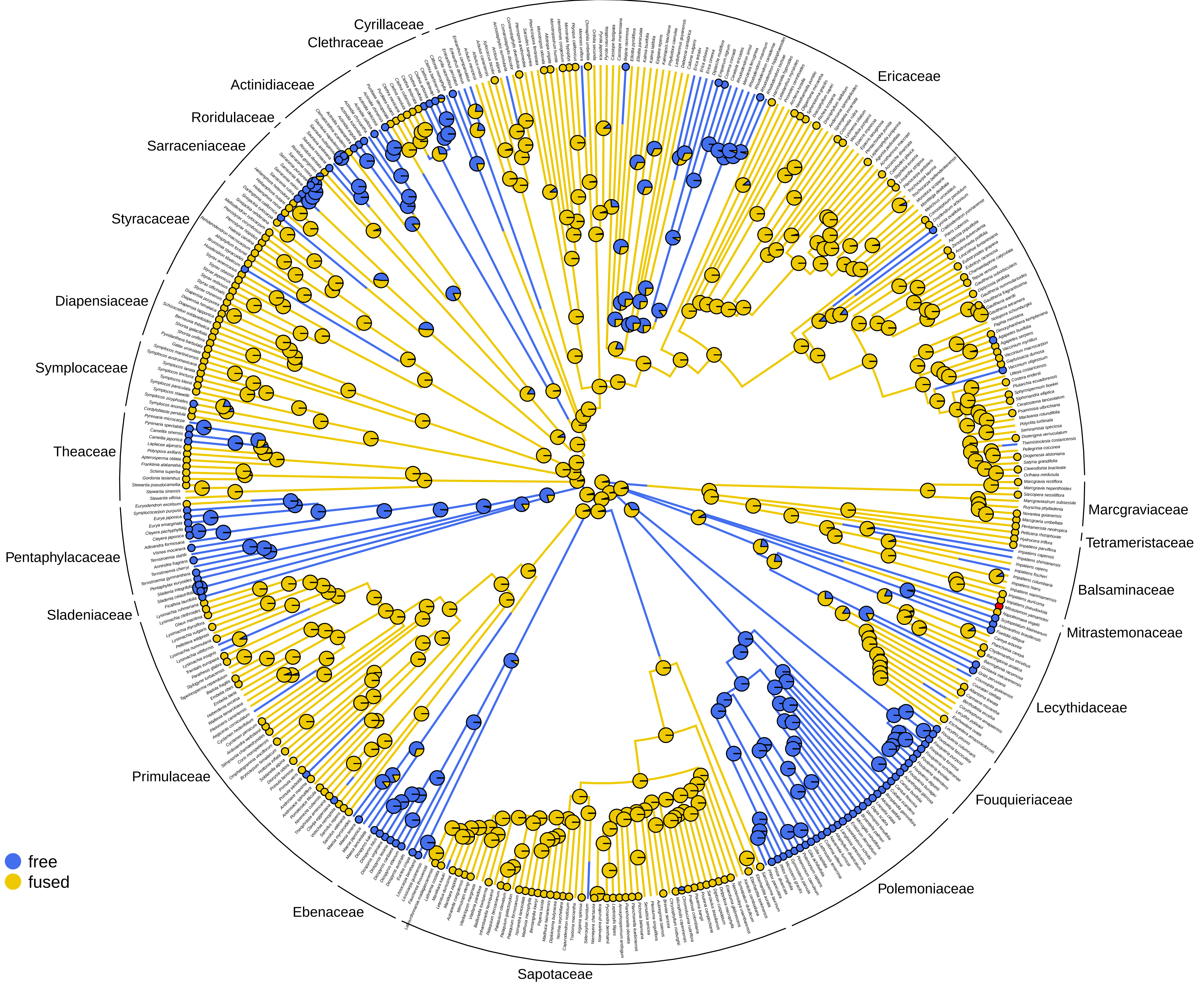
